# MLMarker: A machine learning framework for tissue inference and biomarker discovery

**DOI:** 10.1101/2025.06.10.658642

**Authors:** Tine Claeys, Sam van Puyenbroeck, Kris Gevaert, Lennart Martens

**Affiliations:** VIB-UGent Center for Medical Biotechnology, VIB, Ghent, Belgium; Department of Biomolecular Medicine, Ghent University, Ghent, Belgium; BioOrganic Mass Spectrometry Laboratory (LSMBO), IPHC UMR 7178, University of Strasbourg, CNRS, Strasbourg, 67000, France

**Keywords:** Tissue prediction, Mass spectrometry-based proteomics, Machine learning, Public data reuse, AI

## Abstract

MLMarker is a machine learning tool that computes continuous tissue similarity scores for proteomics data, addressing the challenge of interpreting complex or sparse datasets. Trained on 34 healthy tissues, its Random Forest model generates probabilistic predictions with SHAP-based protein-level explanations. A penalty factor corrects for missing proteins, improving robustness for low-coverage samples. Across three public datasets, MLMarker revealed brain-like signatures in cerebral melanoma metastases, achieved high accuracy in a pan-cancer cohort, and identified brain and pituitary origins in biofluids. MLMarker provides an interpretable framework for tissue inference and hypothesis generation, available as a Python package and Streamlit app.

## Background

Mass spectrometry (MS)-based proteomics has become a cornerstone for protein expression analysis in biological systems. Recent technological advancements in sample preparation^1–3^, instrumentation^4–6^, and computational methods^7–10^ have drastically increased sensitivity, throughput and coverage, enabling the detection and quantification of thousands of proteins across diverse biological samples, all while accommodating ever decreasing amounts of sample. However, as the scale and speed of proteomic dataset generation keeps growing, drawing meaningful biological conclusions remains a challenge due to the inherent complexity and vastness of the proteome^11,12^.

Many downstream interpretation methods rely primarily on the detection of differentially expressed proteins, and these approaches often fall short of providing a holistic, multi-layered comparative analysis. Traditional approaches, such as pairwise differential expression analysis (DEA), aim to narrow the scope of data interpretation by focusing on predefined group comparisons. While these methods offer statistical rigor, they cannot reveal more complex, non-obvious relationships within the data. Additionally, they can be significantly influenced by preprocessing steps such as quantification methods, normalization, and missing value imputation, resulting in a combinatorial explosion of possible outcomes, in turn leading to inconsistent results^13,14^. Capturing the full differential landscape of a sample while considering its broader context, thus remains a significant challenge. The field therefore stands to gain substantially from the development of innovative approaches that can more comprehensively address the data’s complexity.

Interestingly, the large amounts of data being deposited into ProteomeXchange^15^ repositories such as PRIDE^16^ and MassIVE^16^ provide the potential to improve these analysis methods. Indeed, this treasure trove of proteomics data has already led to the development of essential community knowledgebases such as PeptideAtlas^17^, ProteomicsDB^18^, and Scop3P^19^. Furthermore, it has enabled technical advances such as the development of machine learning models like MS2PIP^20^ and DeepLC^21^ that allow greatly improved MS data interpretation through rescoring approaches^7,22^. While the reuse of public data for biological discovery has so far remained limited, it has demonstrated potential in identifying and validating biomarkers^23^, and in conducting meta-analyses on post-translational modifications^24,25^.

Here, we introduce another way of making use of this vast public proteomics data with MLMarker, a machine learning (ML) based tool based on our previously developed tissue prediction model^26^. The basic ML framework of MLMarker was extended to be adaptable and explainable. The tool is designed to complement proteome analysis by using its ML model to compare each sample to a reference atlas of healthy human tissue proteomes applicable to even samples that do not resemble the original training data. Unlike conventional approaches that rely on rigid, pre-assigned group comparisons that typically all originate in the same study, MLMarker provides a continuous, probabilistic measure of tissue similarity across multiple tissues simultaneously. Furthermore, due to its simplistic nature the results are highly explainable. This enables a more nuanced view of sample classification, revealing unexpected tissue relationships that may indicate disease adaptations and/or uncover novel biological phenomena. Importantly, MLMarker thus shifts tissue prediction from a simple classification task to a powerful framework for hypothesis generation, as samples with unexpected tissue similarities can point to new biological mechanisms or disease processes ^27–29^.

We first showcase the newest technical developments followed by a demonstration of the model’s biological applicability on three public datasets: (i) a reanalysis of cerebral melanoma metastases from Zila *et al*. (PXD007592)^30^, (ii) a reanalysis of cerebrospinal fluid from Yamana *et al*. (PXD008029)^31^, and (iii) a reanalysis of the pan-cancer FFPE tissue dataset from Tüshaus *et al*. (MSV000095036)^32^. Together these three case studies illustrate the versatility of MLMarker, showing its potential not only in understanding cancer metastases, but also in analysing biofluid samples and mapping tissue identities in complex multi-cancer datasets. By leveraging a tissue-centric approach, MLMarker offers a new perspective on proteomics data, complementing existing methods and expanding the scope of biological discovery.

MLMarker is available as a pip installable python package and a graphical user interface through the streamlit framework at https://mlmarker.streamlit.app.

## Results

### Adaptability and explainability as core-developments

The initial tissue classification model classifier was developed as a trained Random Forest model that assigns tissue labels from a fixed feature space, trained on high abundant proteins of public proteomics data encompassing healthy tissues acquired with data-dependent acquisition. Although this approach captures complex relationships in high dimensional proteomics data, a standalone estimator is difficult to deploy across heterogeneous due to three recurrent sources of heterogeneity including (i) protein identity; (ii) quantification scale; and (iii) incomplete feature coverage. MLMarker therefore generalises the original model into a Python package that explicitly addresses these issues through three core developments.

First, MLMarker replaces purely probability-based outputs with a Shapley Additive exPlanations (SHAP) based reconstruction of tissue scores, decomposing tissue similarity into per protein contributions. This enables the identification of proteins driving similarity toward, or away from, a given tissue allowing full explainability of the predictions.

Second, MLMarker automatically applies per sample min-max normalization prior to inference. By focusing on within sample ratios, the model becomes less sensitive to global scaling differences introduced by experimental and computational pipelines, becoming applicable to modalities differing from the original training setting (e.g. data-independent acquisition, affinity-based proteomics methods). Third, to accommodate sparse protein coverage, intrinsic to proteomics, the package uses an adjustable penalty factor that modulates how absent proteins affect the SHAP explanation and resulting tissue ranking. The penalty is calibrated using a zero sample SHAP reference computed at initialisation, which represents the model behaviour in the absence of detected proteins and provides a pragmatic quality control anchor for low coverage samples.

These software and methodological extensions transform MLMarker from a static classifier into an adaptable and explainable framework that supports hypothesis generation by linking tissue similarity patterns to the underlying protein evidence and by enabling contextualisation through external knowledge bases and over-representation analyses.

#### Penalty factor

Missing values are intrinsic to mass spectrometry-based proteomics and can arise through distinct mechanisms from both technical and biological origin. These missing values impact MLMarkers ability for accurate prediction as inference is performed in a fixed feature space of 5,979 protein defined during model training. When a protein is not reported within a new sample, its abundance is treated as zero. This creates a systematic bias: samples with low proteome coverage have many features artificially set to zero, which can shift predictions toward tissues that naturally express fewer proteins. To quantify how this ambiguity affects tissue similarity predictions, and to assess whether the introduction of a penalty factor (λ) for handling of missing features improves robustness, we performed controlled missingness simulations. The simulation design, missingness mechanisms, penalty strategies, and evaluation metrics are described in detail in the Methods section.

We focused on two public datasets that approximate the extremes encountered in tissue prediction. A dense liver tissue proteome was obtained from Davis et al.^33^ (PXD009021, target tissue: Liver) and a sparse cerebrospinal fluid from Yamana *et al*. ^34^ (PXD008029, target tissue: Pituitary gland). For each dataset, we started from reprocessed data using ionbot^35^ for peptide identification, FlashLFQ^36^ for quantification. Next, we simulated increasing levels of protein dropout across three different scenarios of missingness: (i) Missing Completely At Random (MCAR), representing technical variation; (ii) Missing Not At Random (MNAR), where low-abundance proteins are preferentially lost, mimicking the detection limits inherent to mass spectrometry; and (iii) Targeted missingness, simulating the loss of tissue-specific marker proteins due to biological differences between sample and reference.

The penalty factor λ is implemented by first computing a zero baseline SHAP profile, and then, for proteins that are absent in the input sample but still contribute nonzero SHAP mass, subtracting λ times their corresponding zero baseline SHAP contribution from the sample specific SHAP value. This method attenuates spurious evidence arising purely from feature absence rather than observed signal, as further specified in the Methods. We evaluated two categories of λ: fixed and adaptive. In the fixed setup, a constant λ was applied across all samples ranging from 0 (no penalization) to 2 (aggressive penalization). In the adaptive penalty setup, λ was set as a function of sample feature coverage in either a linear or logistic function so that penalization increased when coverage decreased.

#### Performance Across Sample Types: Dense vs. Sparse

Our simulations demonstrate context-dependent behavior that informs practical recommendations on when exactly to enable this MLMarker feature.

Looking at the top-1 accuracy e.g.‘is the highest predicted tissue correct?’ (Fig 1a), we observe a generally high accuracy in dense tissues for both MCAR and MNAR scenarios, independent of λ. This is not the case for the spare biofluid sample where the absence of a penalty factor leads inevitably to an extreme drop in performance which is restored by the usage of any sort of λ. When tissue-specific markers were preferentially removed, hence simulating biological divergence, both dense and sparse samples show a dramatic accuracy drop that could be partially rescued by λ. Comparing penalty methods across all dropout rates and missingness methods (Fig 1b) show that λ=1 and the adaptive piecewise λ show the highest performance for dense tissue (95.6% accuracy) and biofluid (77.0% accuracy) respectively.

**Fig 1.**
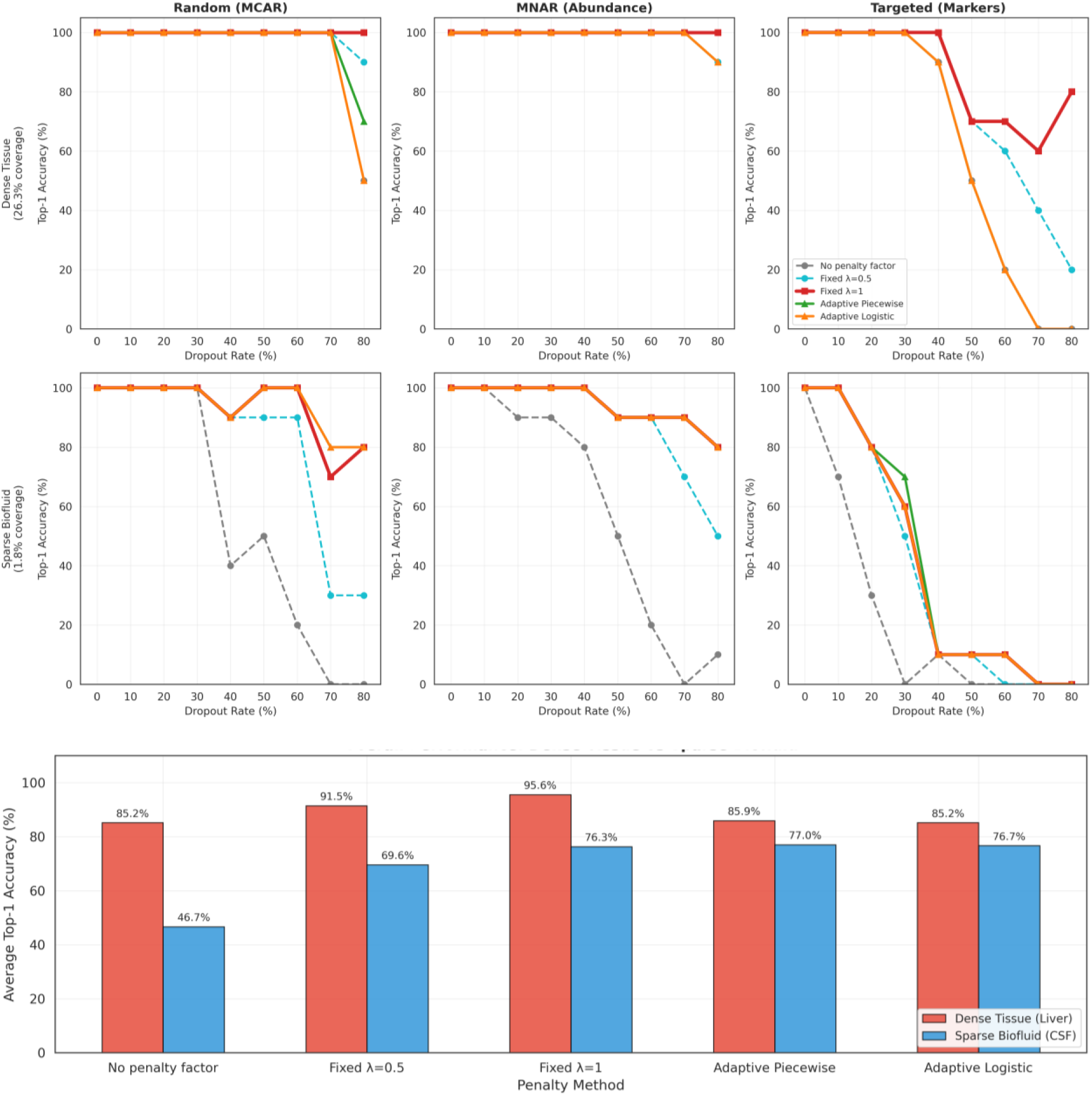
A) Overview of top-1 accuracy (y-axis, is the highest predicted tissue correct?) across missingness patterns (columns), dropout rates (x-axis) and penalty factors (λ) (curve colors) for dense tissue (top) and sparse biofluid (bottom). B) Average Top-1 accuracy across all dropout rates per penalty method for both dense tissue (red) and sparse biofluid (blue)

Top-1 accuracy captures only whether the expected tissue is ranked first, but does not quantify how strongly predictions degrade when the correct tissue drops to lower ranks. We therefore examined rank degradation under targeted missingness, which produced the largest divergence between penalty strategies, and observed that a λ = 1 yielded the most stable rankings in both dense tissue and sparse biofluids (Fig. 2; Supplementary figure 1).

**Fig 2.**
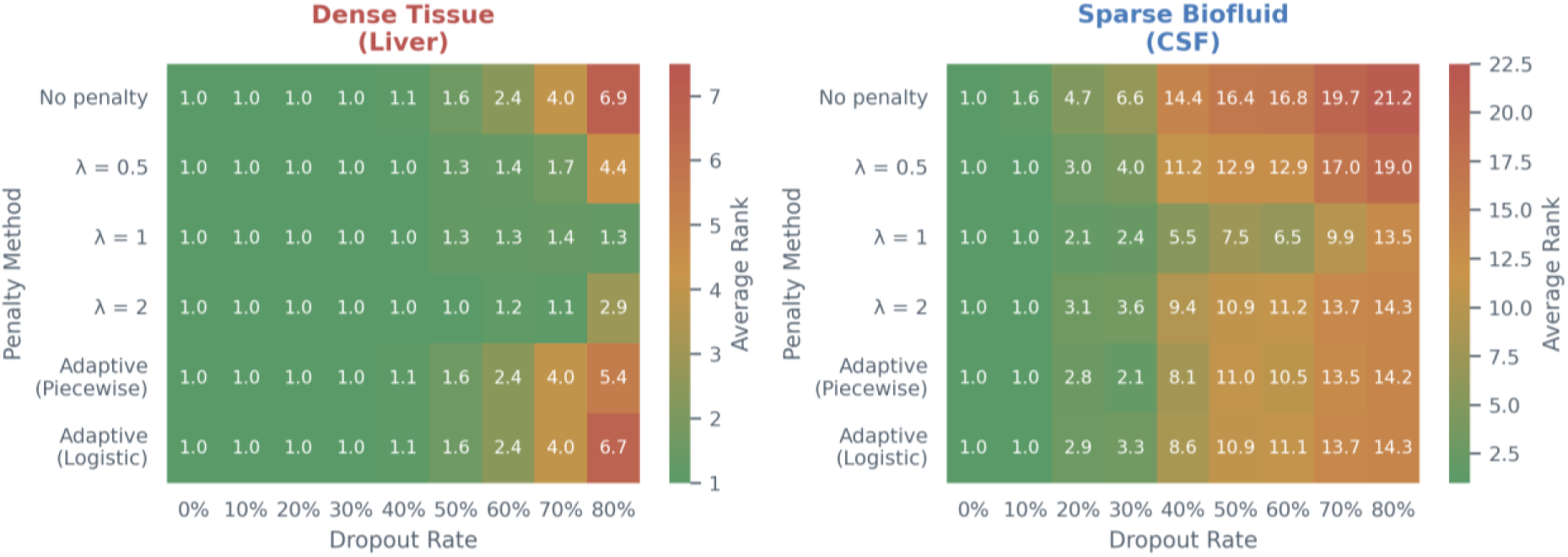
Average rank degradation under targeted missingness across penalty strategies for dense tissue (liver, left) and sparse biofluid (CSF, right). Each heatmap displays the average rank of the correct tissue prediction across increasing dropout rates (x-axis, 0–80%) for six penalty methods (y-axis): no penalty, fixed λ values (0.5, 1, 2), and adaptive strategies (piecewise, logistic). In both samples, λ = 1 consistently maintains low rank values across all dropout levels, outperforming other methods especially under high missingness

With correcting for missing proteins, the risk of overcorrection cannot be left unexplored. To assess this possibility, we compared the accuracy of λ = 1 to λ = 0 and defined overcorrection as any reduction in accuracy relative to the no penalty baseline (Fig 3). In the sparse CSF, λ = 1 improved accuracy consistently across all missingness types and dropout levels, with mean gains of 0.72% under MCAR, 0.62% under MNAR, and 1.06% under targeted missingness. The absence of overcorrection for sparse samples, indicates that down weighting absent features is uniformly advantageous when baseline coverage is low. In the dense liver tissue, the effect depended on the drop-out rate. Under MCAR and MNAR, λ = 1 induced mild overcorrection at low dropout, with mean effects of - 0.28% and - 0.40%, and became beneficial only under extreme information loss above roughly 65 to 70% dropout. In contrast, when missingness preferentially removed targeted informative marker proteins, λ = 1 remained consistently beneficial even in dense tissue, with a mean gain of 1.3%.

**Fig 3.**
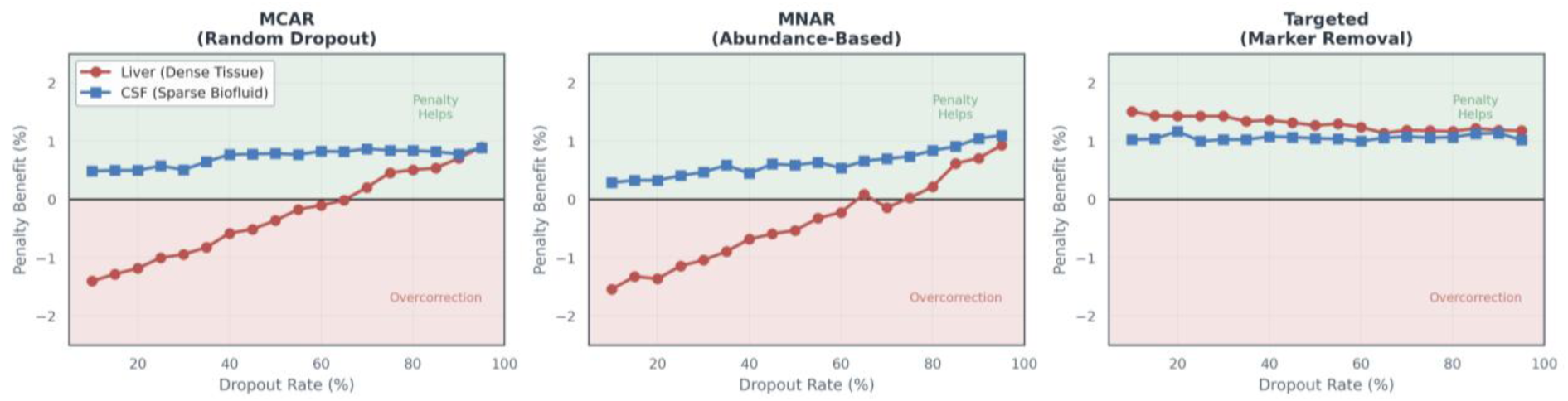
Penalty benefit across dropout conditions for dense tissue (liver) and sparse biofluid (CSF). Each panel shows the change in top-1 accuracy when applying a penalty factor (λ = 1) relative to no penalty (λ = 0) across increasing dropout rates under three missingness mechanisms: MCAR (left, random dropout), MNAR (middle, abundance-based dropout), and targeted removal of marker proteins (right). Positive values indicate improved accuracy (“Penalty helps”), while negative values indicate overcorrection. In CSF, λ = 1 consistently improves accuracy across all dropout levels and missingness types. In liver, λ = 1 causes mild overcorrection at low dropout under MCAR and MNAR but becomes beneficial under high dropout and remains advantageous under targeted missingness.

### Reprocessed data from cerebral melanoma (PXD007592)

Zila *et al*. analysed proteomic profiles of cerebral melanoma metastases to grasp the difference between good and poor treatment response to MAP kinase inhibitors (MAPKi). The samples were obtained through surgical excision, and response state to MAPKi treatment was determined through progression-free survival (PFS). Good responders had a PFS ≥ 6 months and low disease progression, while poor responders had a PFS < 3 months and fast disease progression. The authors observed small molecular differences between good and poor responders, with poor responders exhibiting an epithelial-mesenchymal transition (EMT) signature and drug resistance mechanisms, while good responders showed higher immune activation. To note is that this different response to treatment was determined on a phenotypic level and not on the proteotype of the tumour. Ultimately, the authors associated good response patients with proliferative melanoma tumours and poor response patients with invasive melanoma tumours, but, at 1% false discovery rate (FDR) and a Perseus s₀ of 0.5, could only identify fourteen differentially expressed proteins between the two response states^30^.

This dataset was selected to explore whether cerebral melanoma metastases are classified by MLMarker as brain, reflecting their site of growth, or as skin, reflecting their tissue of origin. As metastatic tumours may retain features of both origins, they were expected to provide a useful model for testing if and how MLMarker resolves their origin. As mentioned, the original analysis identified only minor molecular differences between good and poor responders, with limited separation in unsupervised clustering, suggesting that relevant biological structures remained hidden. By applying MLMarker, we aimed to determine whether tissue similarity scores could uncover this latent structure and offer complementary insight beyond treatment response.

Reprocessing of the dataset was performed using ionbot^35^ for peptide identification, FlashLFQ^36^ for quantification, and MSqRob2^37^ for differential expression analysis. Twenty proteins were found significantly differentially expressed (adjusted p-value < 0.01) and this low variance between response states is also reflected in the UMAP clustering of protein expression values which shows no clear separation between response states (Fig 4A). Only one protein was indicated as differentially expressed in both workflows: the eukaryotic translation initiation factor 3 subunit I (See Supplementary Table S1). Hierarchical clustering identified two groups. These groups are however not very separated in the UMAP space and DEA revealed only 1 differentially expressed protein (Suypplementary figure 2). Even beside the fact that this approach is statistically incorrect as it uses the same data twice, violating statistical independence, and is often referred to as double-dipping^38^, it is also an indication of the homogenous nature of the melanoma samples.

**Fig 4.**
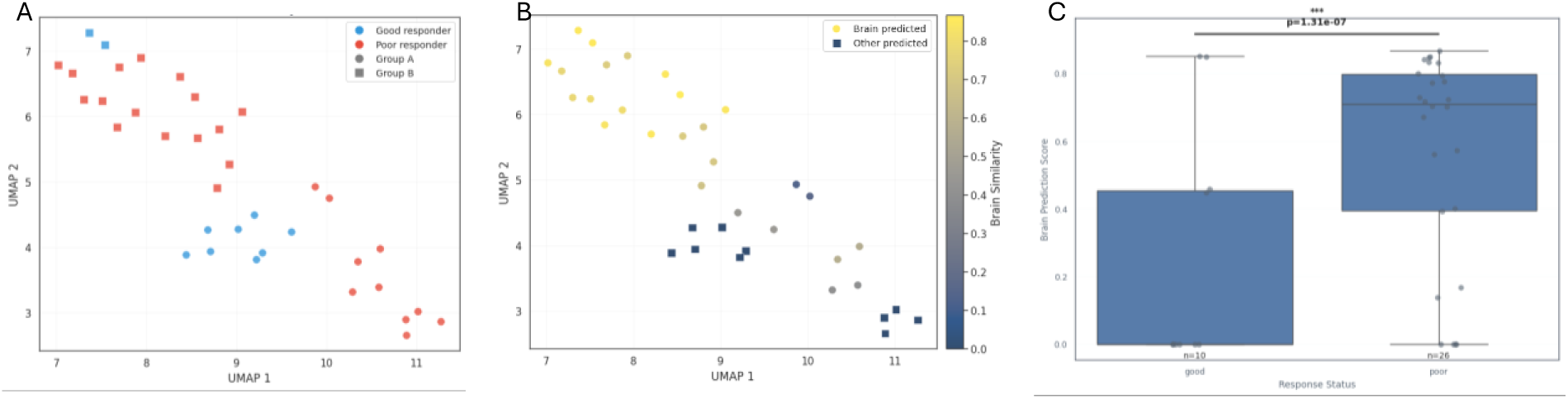
A) UMAP embedding of protein expression values (x- and y-axes) coloured by response status and clustering groups. No separation is observed, consistent with minimal differential expression. B) UMAP embedding coloured by MLMarker-derived brain similarity scores, revealing a clear gradient not captured by expression-based clustering. C) Brain similarity scores (y-axis) for good (n = 10) and poor (n = 26) responders (x-axis). Poor responders show significantly higher brain similarity (Mann–Whitney p = 1.31 × 10⁻⁷).

Here MLMarker presents orthogonal information. When using the same UMAP of protein expression values but using colour to indicate predicted brain similarity with MLMarker, this shows a gradient of brain similarity (Fig 4B). Furthermore, the brain prediction state significantly correlates with response state. Indeed, poor responders had significantly higher brain similarity scores than good responders (Mann-Whitney U test, p = 1.31E^-7^, Fig 4C).

This resulted in four subpopulations: poor responders with high brain similarity (twenty samples) and with low brain similarity (six samples), and good responders with high brain similarity (four samples) and low similarity (six samples). Notably, when samples were not predicted as brain, their highest-scoring alternative tissues were either lung or monocytes.

To determine whether MLMarker’s predictions were influenced by data quality or technical artifacts, we analysed (i) total protein intensity across samples, (ii) the intensity of the MLMarker features and (iii) the general feature coverage. Indeed, MLMarker relies on a set 5,979 of predefined proteins seen during training, meaning it can only process and generate predictions using the proteins in this feature set. Any missing proteins are assigned a value of zero, potentially affecting the accuracy and reliability of the predictions.

Total protein intensity showed a significant correlation with brain cluster assignment, a correlation that remained when only using the MLMarker features (Fig 5A and 5B, p=0.012 and p=0.02 respectively). However, samples predicted as brain and not as brain did not have a significantly different distribution in the MLMarker feature coverage (Fig 5C, p=0737). To further explore if prediction is hence driven by that intensity, or by a different population of MLMarker features present, we studied the intensity distribution between features and non-features. This demonstrated that the MLMarker features in brain samples have a higher intensity but importantly, this proportionally accounts for 56.7% of total intensity, which is the same signal proportion as in the samples not predicted as brain (58.3%, Fig 5D). Moving towards a sample-level analyses, a near-perfect correlation between the intensity of MLMarker features vs not-MLMarker features can be observed (R=0.98, Fig 5E). This all comes to the conclusion that the observed intensity variation is a global sample-level effect unrelated to MLMarker’s feature space. Together, these results demonstrate that MLMarker’s brain predictions are not driven by data quality, quantity, or intensity artifacts.

**Fig 5.**
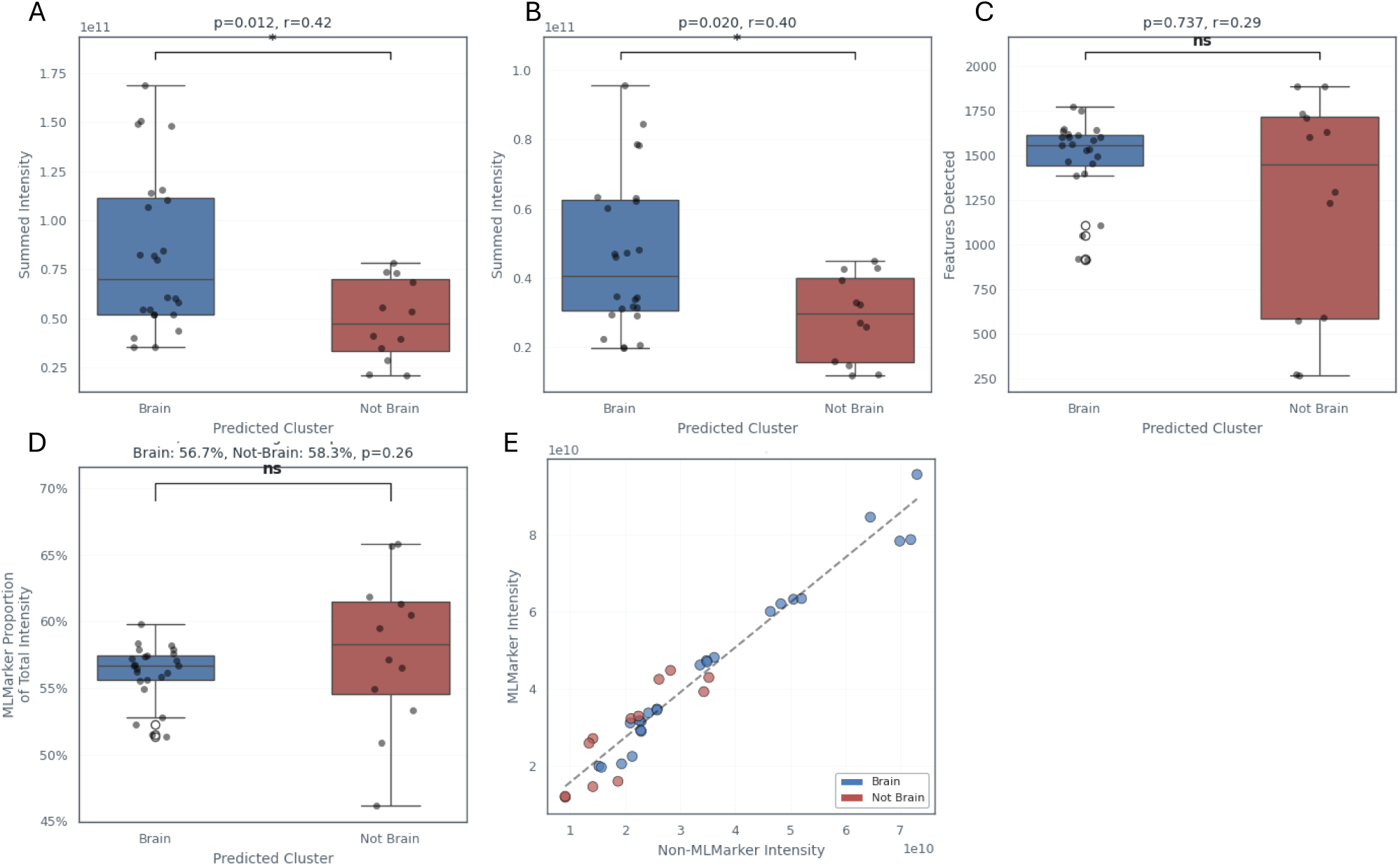
A) Total protein intensity (y-axis) compared between samples predicted as brain and not-brain (x-axis). Brain-predicted samples show significantly higher total intensity (p = 0.012). B) Summed intensity of MLMarker feature proteins (y-axis) for brain vs. not-brain samples (x-axis), also significantly higher in brain-predicted samples (p = 0.020). C) Number of detected MLMarker features (y-axis) across brain vs. not-brain samples (x-axis), showing no significant difference (p = 0.737). D) Proportion of total sample intensity attributable to MLMarker features (y-axis) for brain vs. not-brain samples (x-axis). Both groups show similar proportions (56.7% vs. 58.3%, p = 0.26), indicating no enrichment of MLMarker features.

To further investigate the tumour profiles, we took two distinct approaches. First, we utilized the insights from the MLMarker model by SHAP values. A negative SHAP value indicates that a protein negatively influences the predicted tissue classification, while a positive SHAP value suggests a positive impact. It is also important to emphasize that the absence of proteins can be informative in this context, and the non-detection of proteins can thus influence the prediction process. Second, we applied the tissue classifications generated by MLMarker to define groups for DEA with MSqRob2^37^, comparing brain against non-brain samples.

#### SHAP analysis of MLMarker decision

To understand the model’s decision process, we plotted the SHAP values across the four subpopulations (Fig 6). These values quantify the contribution of each feature to the model’s predictions. Notably, brain-predicted samples exhibit significantly higher SHAP values (p= 6.73 × 10⁻¹⁶), suggesting that several proteins positively influence the model’s decision-making. In contrast, non-brain-predicted samples show lower SHAP values, indicating that the model relies less on tissue-specific markers for brain classification. Instead, classification appears driven by a broader proteomic profile, potentially incorporating less tissue-specific features. This reinforces the conclusions from the previous analysis, stating that the classification is driven by a diverse population of proteins and not by abundances.

**Fig 6.**
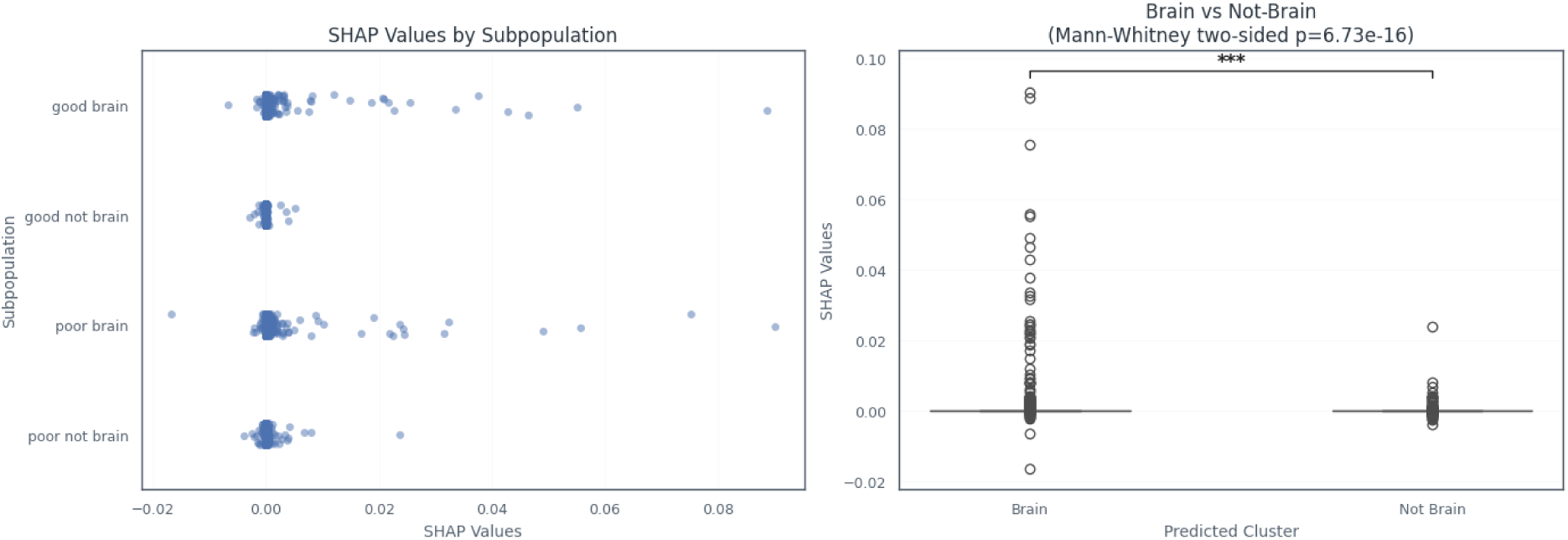
A) SHAP values (x-axis) across the four subpopulations (y-axis: good-brain, good-not-brain, poor-brain, poor-not-brain). Brain-predicted samples show consistently higher SHAP values, indicating stronger positive feature contributions to the model’s decision. B) Distribution of SHAP values (y-axis) for samples predicted as brain vs. not-brain (x-axis). Brain-predicted samples exhibit significantly higher SHAP values (Mann–Whitney two-sided p = 6.73 × 10⁻¹⁶), demonstrating that MLMarker relies on a broader set of positively contributing proteins when assigning brain similarity.

To gain a more in-depth understanding of decision-driving proteins in each subpopulation, we selected the top 1% highest-ranking SHAP value proteins per subpopulation for an over-representation analysis, revealing a clear tissue-specific signature (Supplementary figure 3). Unsurprisingly, brain-predicted samples, regardless of response state were enriched for brain-associated processes. In contrast, non-brain predicted samples showed a sharp decline in brain-related functions among poor responders and a complete absence in good responders. Instead, good responders exhibit an increase in immune-related processes, which are entirely absent in the other groups.

#### Differential expression analysis with MSqRob

To fully demonstrate MLMarker’s potential as a hypothesis generator, we performed two types of DEA. As mentioned earlier, comparison of responders and non-responders, identified twenty differentially expressed proteins (DEPs) with an adjusted p-value < 0.01, a similar number of DEPs compared to the original results. However, when we performed the same analysis between brain-predicted and non-brain-predicted samples, we observed a high increase in DEPs, totalling 255 proteins. Of these, 217 were upregulated within brain-predicted samples, and 38 were upregulated within not brain-predicted samples. These results far exceed the number of DEPs identified in the original responder comparison (Fig 7).

**Fig 7:**
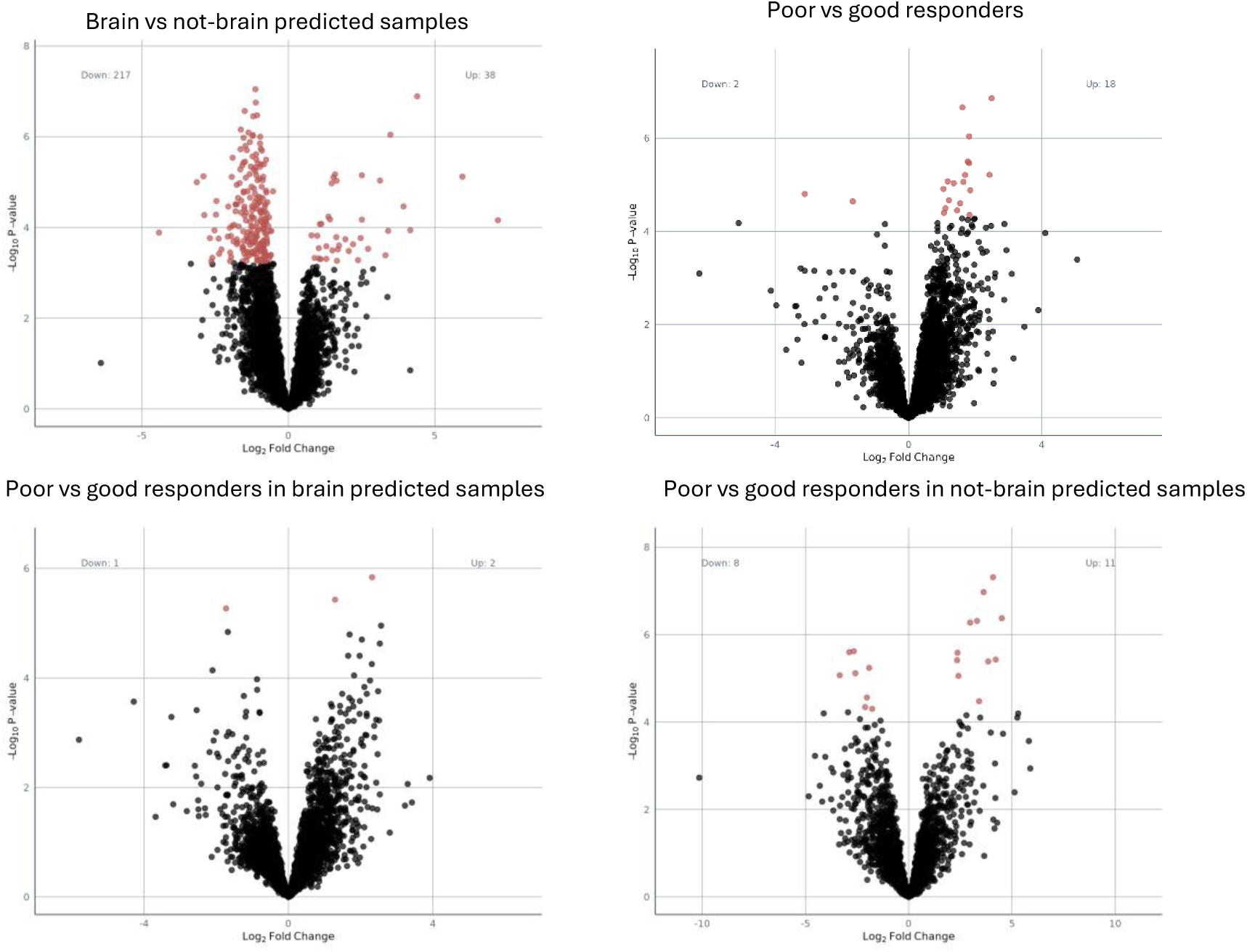
Differential expression analysis volcano plots with log fold change on X-axis, and -log_10_ adjusted p-value following Benjamini-Hochberg correction on Y-axis. Red dots indicate proteins with adjusted p-value < 0.01.Top panel: Comparison of brain-predicted vs. non-brain-predicted samples (left) identified 255 DEPs (adjusted p-value < 0.01), a tenfold increase over the responder vs. non-responder comparison (right).Bottom panel: Within brain-predicted (left) and non-brain-predicted (right) subpopulations DEA between good and poor responders, 3 DEPs were identified in brain-predicted samples, while 19 DEPs were found in non-brain-predicted samples.

Next, we performed DEA between good and poor responders within the brain-predicted and non-brain-predicted subpopulations separately. This analysis revealed three DEPs in brain-predicted samples, while 19 DEPs were identified in non-brain-predicted samples (Fig 7). Taken together, these subgroup-dependent results support the use of MLMarker classification as a stratification variable for downstream analyses and hypothesis generation, as it reveals proteomic structures that might not be apparent in the initial comparison and can yield candidate proteins for follow-up.

#### Further analysis of DEPs in brain vs non-brain

Analysing the 255 DEPs (Table S1), a striking observation is their limited brain specificity. Indeed, over-representation analysis reveals two distinct molecular signatures (Supplementary figure 4) Proteins overexpressed in brain-predicted samples include some that are involved in neuronal processes, yet their functions extend beyond the brain. Instead, they are associated with metabolism, extracellular space, secretion, protein binding, and enzyme activity. Additionally, these proteins are linked to metabolic pathways such as the TCA cycle, and Parkinson’s disease. Notably, the molecular functions “cadherin binding” and “heat shock protein binding” emerge as enriched, both of which were previously associated with an invasive tumour phenotype^30^.

In contrast, the 31 proteins overexpressed in non-brain-predicted samples are predominantly linked to protein activation cascades, blood coagulation, and extracellular processes. From a published list of 76 known protein markers of tumour invasion^39^, ten were found among the DEPs in brain-predicted samples, while none were present in the non-brain predicted DEPs. These ten proteins are primarily involved in metabolism, cytoskeleton organization and signalling (Table 1).

**Table 1:**
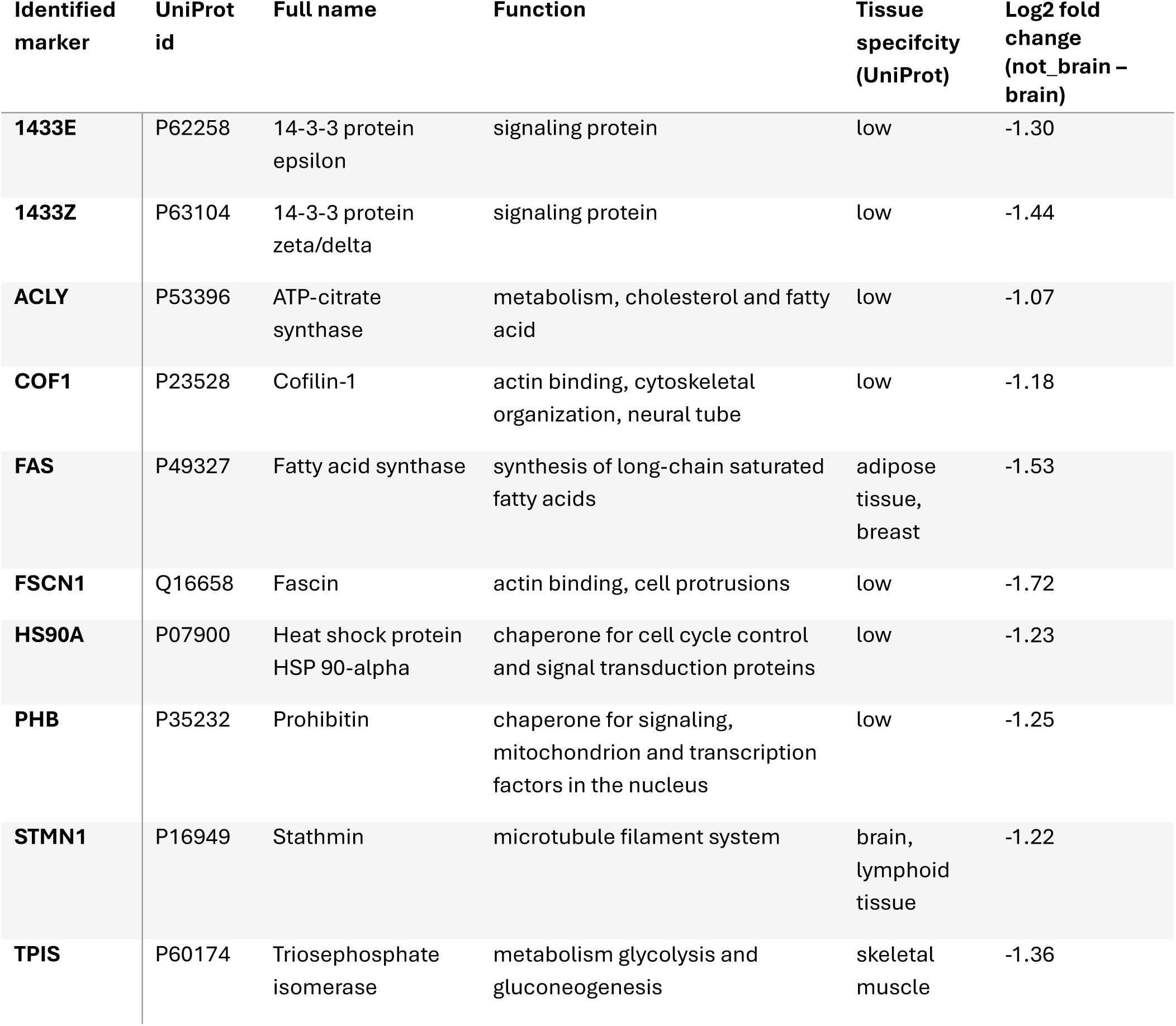
Differentially expressed proteins that are known markers of tumour invasion according to Pouliquen et al. ^39^

#### Circularity and double-dipping

Double-dipping mainly occurs when the same dataset is used to define a contrast and then used to demonstrate the significance of that contrast. This is not expected within the MLMarker approach as MLMarker operates on the level of individual samples and produces tissue similarities by comparing to an external reference atlas, rather than learning any separation between sample groups. In this way, MLMarker primarily functions as an annotation layer and does not include sample variation. However, it is worth checking this empirically as both MLMarker and MSqRob are applied to the same quantitative protein measurements, and proteins that drive tissue similarity in MLMarker could also be among those that show differential regulation between the MLMarker defined groups. This coupling would not imply methodological invalidity, but it could bias interpretation if the MLMarker feature space were systematically enriched for low adjusted p values. We therefore assessed potential circularity by comparing p value distributions and differential expression rates inside and outside the MLMarker feature set, and by examining whether SHAP based importance aligns with statistical evidence beyond what is expected under a simple enrichment effect.

First, we examined the proportion of DEPs in the MLMarker features, followed by a correlation analysis between SHAP values from MLMarker predictions and adjusted p-values from MSqRob when comparing brain and not-brain predicted samples. Comparing the p-value distributions between MLMarker features and not MLMarker features revealed a similar pattern (Fig. 8A). This similarity persisted when restricting to DEPs with log_2_fold changes greater than |1|. Within the MLMarker features the proportion of DEPs (13.6 %) closely matches that within not MLMarker features (11.9%). A Mann-Whitney U test showed no significant difference in the p-value distribution for all proteins (p = 0.08) but indicated a slight skew in more differentially regulated proteins among MLMarker features when considering log_2_ fold change >|1| (p = 0.04).

**Fig 8.**
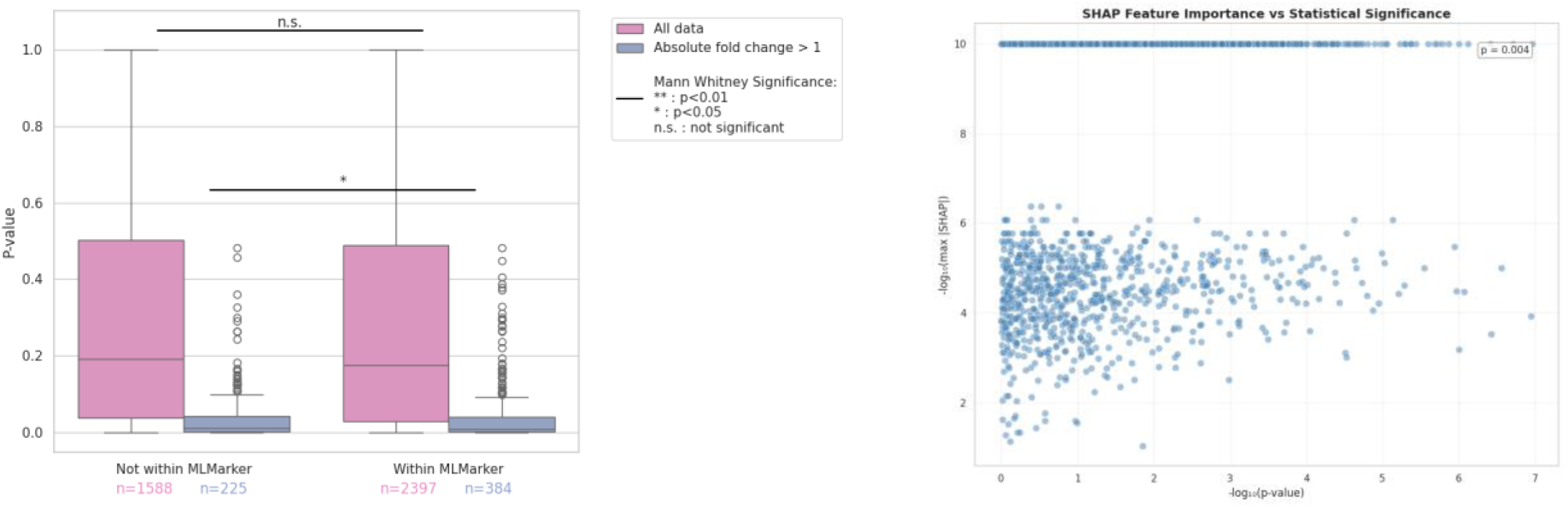
A) Boxplot showing the distribution of p-values for proteins outside (not within) and inside (within) the MLMarker feature space. All proteins, independent of their log_2_ fold change, are shown in green, while proteins with absolute log_2_ fold change > 1 are highlighted in orange. Mann–Whitney U tests indicate no significant difference between groups for all proteins (p = 0.08), while proteins with absolute fold change > 1 show a significant difference (p = 0.04). B) Scatterplot comparing on the Y-axis the log_10_-transformed maximal absolute SHAP value and on the X-axis the minimal - log-transformed raw p-values. Each point represents a protein feature. No significant correlation between SHAP values and adjusted p-values was identified (Spearman ρ = 0.004, p = 0.938).

Next, we evaluated whether the p-value of MLMarker features correlated with the models SHAP values (Fig. 8B). Plotting maximal absolute SHAP values against the p-values from MSqRob, we observed no significant correlation: Spearman ρ = 0.004 (p = 0.938). This indicates that while SHAP-positive proteins may show enriched differential signal as a group as seen in the previous analysis, there is no systematic scaling relationship between feature importance and statistical significance. Overall, the limited correlation indicates partial overlap between SHAP values and differential expression results without evidence of circular analysis.

This aligns with the distinct nature of the measures: SHAP values capture tissue-specific patterns learned from external data, while differential expression reflects cohort-specific statistics. Thus, the results support SHAP-based interpretation as a complementary approach rather than one prone to circular reasoning or data leakage.

### Reprocessed data from cerebrospinal fluid (PXD008029)

In this study (PXD008029^40^), an experimental workflow was developed to maximize protein identifications from human cerebrospinal fluid (CSF) and human plasma. These biofluids underwent depletion of highly abundant proteins and fractionation to enhance the detection of lower-abundance proteins, including potential tissue-leakage markers. Because MLMarker was trained on healthy, solid tissue proteomes and performs inference in a fixed feature space, these biofluids represent a stringent test case for missing feature handling.

We suspected that this sample type posed a challenge for MLMarker as MLMarker was not trained on any biofluids and relies on the previously discussed fixed feature space. Indeed, MLMarker can only use the proteins it was trained upon and requires all proteins to be associated with a quantitative value. To mitigate skewed predictions in sparse datasets, two strategies can be employed. First, the relationship between prediction certainty and protein content can be assessed. The impact of the penalty factor is evident when predicting an empty sample: without correction, the model assigns a 19% tissue similarity score to bone marrow, which drops to 0.5% upon applying the penalty factor (Supplementary figure 5).

When predicting the CSF samples, pituitary gland and brain are the two highest ranking tissues independent of using a penalty factor. However, introducing the penalty factor reduces the similarity score of lower ranking tissue classes such as bone marrow, testis, lung and monocytes. The plasma predictions demonstrate a similar drop in bone marrow. In those samples, adipose tissue is the first ranking tissue followed by monocytes. This can be explained by the lack of any blood-related tissue within the MLMarker training data (Fig 9).

**Fig 9.**
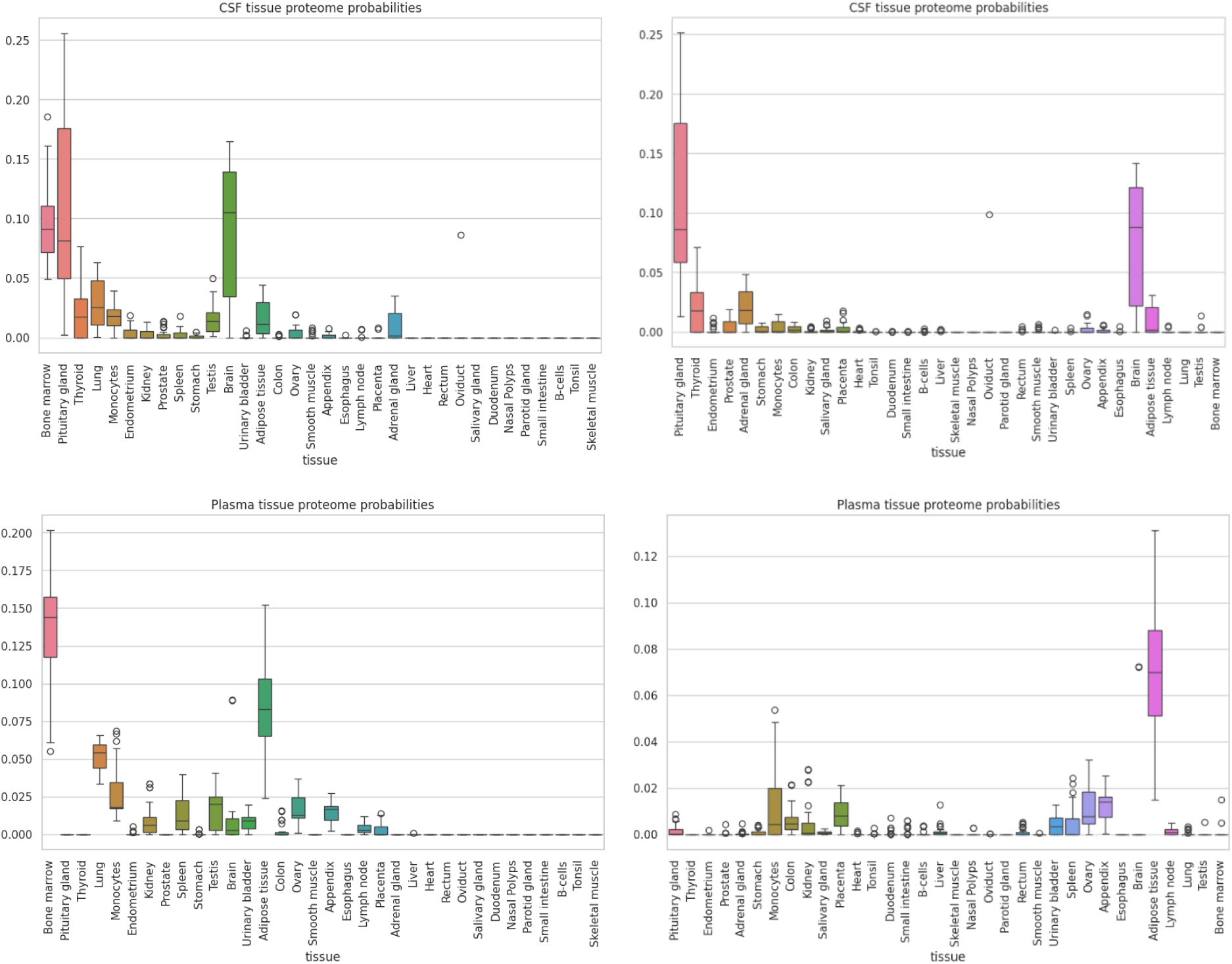
Top panels: CSF tissue similarity before (left) and after (right) penalty factor correction. Bottom panels: Plasma tissue similarity before (left) and after (right) penalty factor correction.

The balance between quantity and prediction confidence is evident in the CSF samples. A strong correlation exists between the number of identified spectra (1% FDR) and the similarity scores of pituitary gland and brain (Pearson correlation of respectively 0.60 and 0.45). However, even low-protein-content samples can yield high-confidence predictions if they contain highly specific proteins (see Supplementary figure 6). For the plasma samples, the correlation is less pronounced for adipose tissue and monocytes (Pearson correlation of respectively 0.23 and 0.44) which can again be explained by the complete absence of any blood-related training data in MLMarker.

#### SHAP analysis of CSF samples within the Streamlit application

MLMarker offers an intuitive and user-friendly interface through a Streamlit application, which is available online and requires no installation. This allows researchers without coding expertise to easily explore the tissue predictions of their samples, making advanced proteomic data analysis accessible to a broader user group. What follows is an example of how MLMarker’s functionalities can be used to gain deeper insights into the proteins driving tissue classifications, facilitated by the interactive features of the Streamlit interface.

The SHAP analysis functionality is visualized through tissue-specific bar plots, which highlight both positively and negatively contributing proteins. For example, in a CSF sample with high predicted probabilities for both the pituitary gland and brain, SHAP values clearly reveal an abundance of proteins reinforcing these tissues, while also showing a substantial number of proteins negatively influencing the bone marrow prediction (Fig 10a).

**Fig 10.**
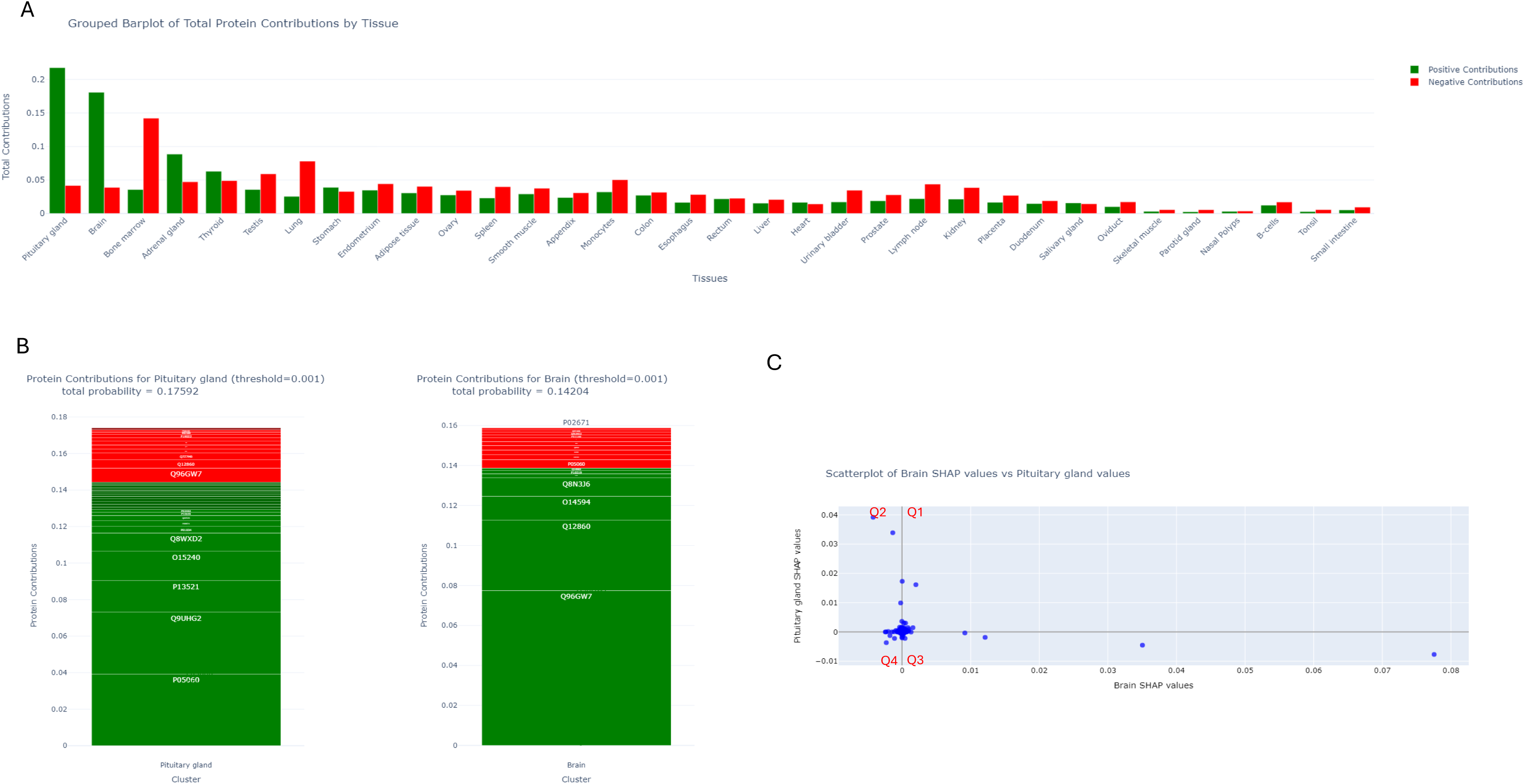
Data from CSF dataset (PXD0080209) A) Sample level grouped bar plot showing positive (green) and negative (red) SHAP contributions per tissue. B) Tissue level bar plot showing positive and negative SHAP contributions for pituitary gland (left) and brain (right). C) Scatterplot of SHAP protein values within brain (x-axis) and pituitary gland (y-axis) showing the interplay of proteins across tissues with quartiles indicated.

Further enhancing the analysis, MLMarker offers an interactive tissue-specific visualization, where bar size reflects the magnitude of each protein’s contribution. This feature allows users to easily browse and identify potentially interesting proteins, enhancing their ability to pinpoint key drivers of tissue classification (Fig 10b). To explore the relationship between protein contributions across tissues, we also generated a scatterplot of adjusted SHAP values for the same CSF sample with high pituitary gland and brain probabilities. This scatterplot reveals how different proteins contribute to each tissue prediction (Fig 10c). (i) those that positively contribute to both tissues, Q1; (ii) those that enhance pituitary gland classification while negatively impacting brain prediction, Q2; (iii) those that strengthen brain classification while reducing pituitary gland tissue similarity score, Q3 and; (iv) those that negatively contribute to both tissues, Q4.

This interactive analysis, seamlessly integrated into the Streamlit application, thus offers an accessible, user-friendly way for researchers to explore and evaluate protein contributions to tissue classifications, making it easier to identify potential biomarkers.

A closer examination of the top tissue-specific contributors confirms their biological relevance. The twenty most influential proteins in the pituitary gland classification predominantly belong to secretory and neuronal pathways, aligning with the known function of this tissue. Likewise, the key proteins driving brain classification are well-established brain-associated markers, several of which are already known to be present in CSF (Tables 2 and 3).

**Table 2.**
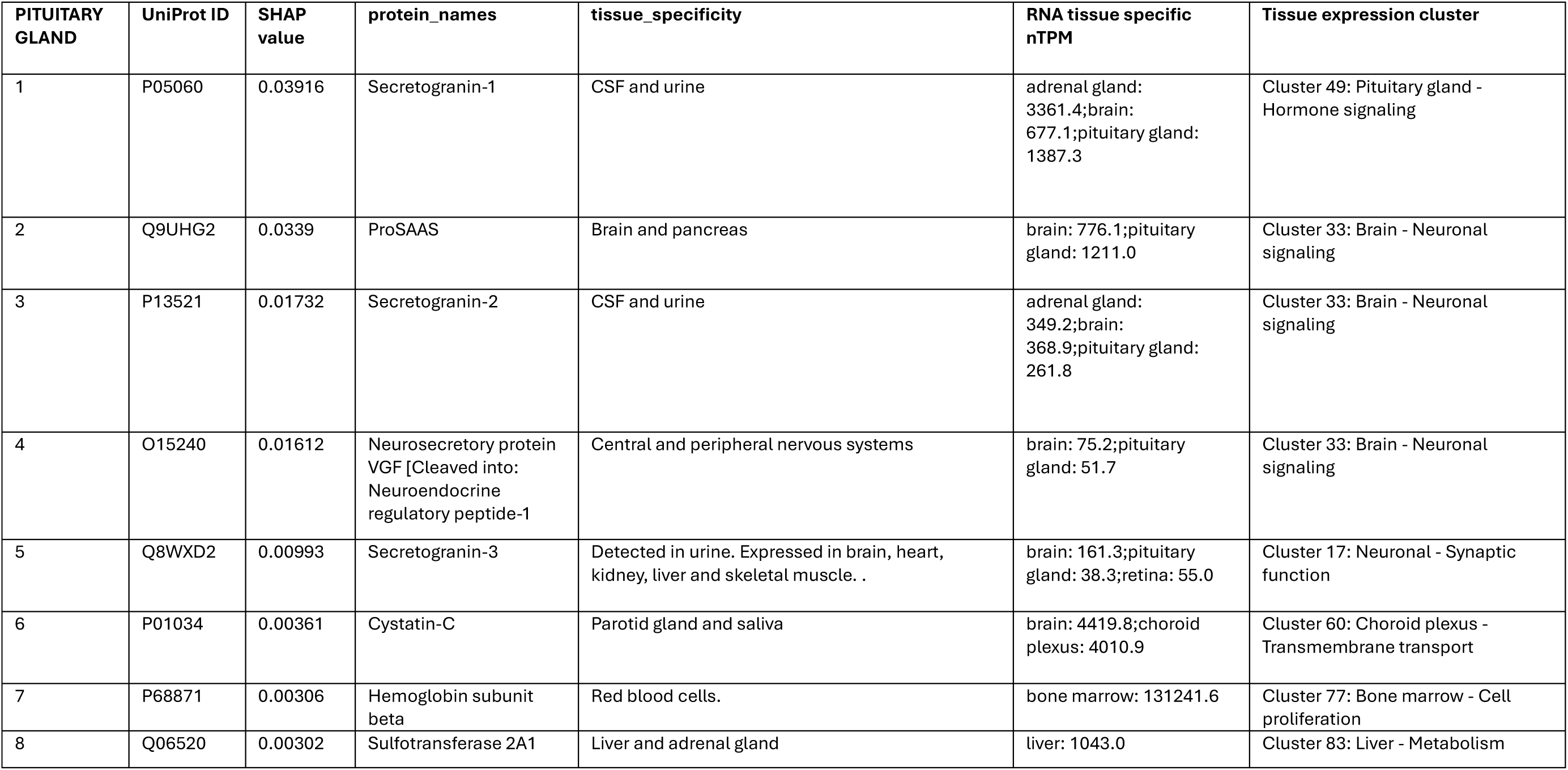
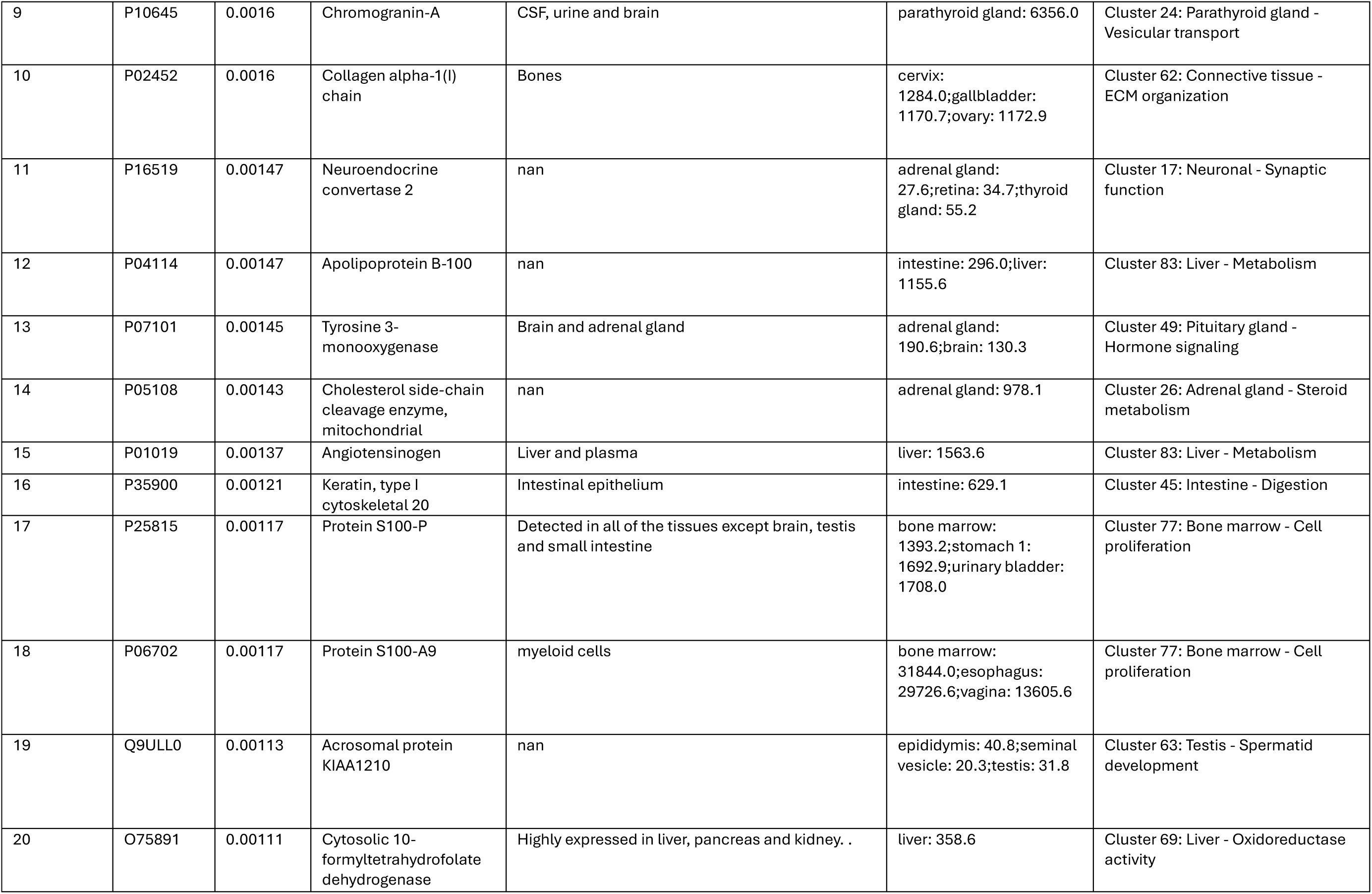
Top 20 proteins positively contributing to pituitary gland tissue prediction, ranked by SHAP value. Gene names, protein names, and tissue specificity annotations were retrieved from UniProt based on the corresponding UniProt identifiers with nan indicating not annotated natively in UniProt. RNA expression levels and tissue-specific expression clusters were obtained from the Human Protein Atlas (HPA). RNA levels are indicated by normalized transcript per million (nTPM) allowing comparison across all HPA samples. Clusters are assigned based on gene to gene distance and shared nearest neighbor clustering to find clusters of genes with similar expression profiles within the graph. Clusters are then manually annotated using overrepresentation analysis including Gene Ontology, KEGG, Reactome, PanglaoDB, TRRUST and HPA classifications.

| PITUITARY GLAND | UniProt ID | SHAP value | protein_names | tissue_specificity | RNA tissue specific nTPM | Tissue expression cluster |
| --- | --- | --- | --- | --- | --- | --- |
| 1 | P05060 | 0.03916 | Secretogranin-1 | CSF and urine | adrenal gland: 3361.4;brain: 677.1;pituitary gland: 1387.3 | Cluster 49: Pituitary gland - Hormone signaling |
| 2 | Q9UHG2 | 0.0339 | ProSAAS | Brain and pancreas | brain: 776.1;pituitary gland: 1211.0 | Cluster 33: Brain - Neuronal signaling |
| 3 | P13521 | 0.01732 | Secretogranin-2 | CSF and urine | adrenal gland: 349.2;brain: 368.9;pituitary gland: 261.8 | Cluster 33: Brain - Neuronal signaling |
| 4 | O15240 | 0.01612 | Neurosecretory protein VGF [Cleaved into: Neuroendocrine regulatory peptide-1 | Central and peripheral nervous systems | brain: 75.2;pituitary gland: 51.7 | Cluster 33: Brain - Neuronal signaling |
| 5 | Q8WXD2 | 0.00993 | Secretogranin-3 | Detected in urine. Expressed in brain, heart, kidney, liver and skeletal muscle. . | brain: 161.3;pituitary gland: 38.3;retina: 55.0 | Cluster 17: Neuronal - Synaptic function |
| 6 | P01034 | 0.00361 | Cystatin-C | Parotid gland and saliva | brain: 4419.8;choroid plexus: 4010.9 | Cluster 60: Choroid plexus - Transmembrane transport |
| 7 | P68871 | 0.00306 | Hemoglobin subunit beta | Red blood cells. | bone marrow: 131241.6 | Cluster 77: Bone marrow - Cell proliferation |
| 8 | Q06520 | 0.00302 | Sulfotransferase 2A1 | Liver and adrenal gland | liver: 1043.0 | Cluster 83: Liver - Metabolism |
| 9 | P10645 | 0.0016 | Chromogranin-A | CSF, urine and brain | parathyroid gland: 6356.0 | Cluster 24: Parathyroid gland - Vesicular transport |
| 10 | P02452 | 0.0016 | Collagen alpha-1(I) chain | Bones | cervix: 1284.0;gallbladder: 1170.7;ovary: 1172.9 | Cluster 62: Connective tissue - ECM organization |
| 11 | P16519 | 0.00147 | Neuroendocrine convertase 2 | nan | adrenal gland: 27.6;retina: 34.7;thyroid gland: 55.2 | Cluster 17: Neuronal - Synaptic function |
| 12 | P04114 | 0.00147 | Apolipoprotein B-100 | nan | intestine: 296.0;liver: 1155.6 | Cluster 83: Liver - Metabolism |
| 13 | P07101 | 0.00145 | Tyrosine 3-monooxygenase | Brain and adrenal gland | adrenal gland: 190.6;brain: 130.3 | Cluster 49: Pituitary gland - Hormone signaling |
| 14 | P05108 | 0.00143 | Cholesterol side-chain cleavage enzyme, mitochondrial | nan | adrenal gland: 978.1 | Cluster 26: Adrenal gland - Steroid metabolism |
| 15 | P01019 | 0.00137 | Angiotensinogen | Liver and plasma | liver: 1563.6 | Cluster 83: Liver - Metabolism |
| 16 | P35900 | 0.00121 | Keratin, type I cytoskeletal 20 | Intestinal epithelium | intestine: 629.1 | Cluster 45: Intestine - Digestion |
| 17 | P25815 | 0.00117 | Protein S100-P | Detected in all of the tissues except brain, testis and small intestine | bone marrow: 1393.2;stomach 1: 1692.9;urinary bladder: 1708.0 | Cluster 77: Bone marrow - Cell proliferation |
| 18 | P06702 | 0.00117 | Protein S100-A9 | myeloid cells | bone marrow: 31844.0;esophagus: 29726.6;vagina: 13605.6 | Cluster 77: Bone marrow - Cell proliferation |
| 19 | Q9ULL0 | 0.00113 | Acrosomal protein KIAA1210 | nan | epididymis: 40.8;seminal vesicle: 20.3;testis: 31.8 | Cluster 63: Testis - Spermatid development |
| 20 | O75891 | 0.00111 | Cytosolic 10-formyltetrahydrofolate dehydrogenase | Highly expressed in liver, pancreas and kidney. . | liver: 358.6 | Cluster 69: Liver - Oxidoreductase activity |

**Table 3.** Top 20 proteins positively contributing to brain tissue prediction, ranked by SHAP value. Gene names, protein names, and tissue specificity annotations were retrieved from UniProt based on the corresponding UniProt identifiers with nan indicating not annotated natively in UniProt. RNA expression levels and tissue-specific expression clusters were obtained from the Human Protein Atlas (HPA). RNA levels are indicated by normalized transcript per million (nTPM) allowing comparison across all HPA samples. Clusters are assigned based on gene to gene distance and shared nearest neighbor clustering to find clusters of genes with similar expression profiles within the graph. Clusters are then manually annotated using overrepresentation analysis including Gene Ontology, KEGG, Reactome, PanglaoDB, TRRUST and HPA classifications.

| BRAIN | UniProt ID | SHAP value | Protein Name | UniProt tissue specificity | HPA RNA tissue specific nTPM | HPA Tissue expression cluster |
| --- | --- | --- | --- | --- | --- | --- |
| 1 | Q96GW7 | 0.07755 | Brevican core protein | Expressed in the retina, CSF and urine | brain: 245.5 | Cluster 33: Brain - Neuronal signaling |
| 2 | Q12860 | 0.03504 | Contactin-1 | Strongly expressed in brain and in neuroblastoma and retinoblastoma cell lines. Lower levels of expression in lung, pancreas, kidney and skeletal muscle. . | brain: 125.0 | Cluster 64: Neuronal - Transcription |
| 3 | O14594 | 0.01211 | Neurocan core protein | Detected in CSF and Brain | brain: 102.1 | Cluster 61: Brain - Neuronal signaling |
| 4 | Q8N3J6 | 0.00916 | Cell adhesion molecule 2 | nan | brain: 50.2;retina: 92.0 | Cluster 17: Neuronal - Synaptic function |
| 5 | O15240 | 0.00202 | Neurosecretory protein VGF [Cleaved into: Neuroendocrine regulatory peptide-1 | Central and peripheral nervous systems | brain: 75.2;pituitary gland: 51.7 | Cluster 33: Brain - Neuronal signaling |
| 6 | P16519 | 0.00159 | Neuroendocrine convertase 2 | nan | adrenal gland: 27.6;retina: 34.7;thyroid gland: 55.2 | Cluster 17: Neuronal - Synaptic function |
| 7 | Q14982 | 0.00132 | Opioid-binding protein/cell adhesion molecule | nan | brain: 36.6;parathyroid gland: 10.0;retina: 11.0 | Cluster 22: Brain - Synaptic signal transduction |
| 8 | P54868 | 0.00099 | Hydroxymethylglutaryl-CoA synthase, mitochondrial | High expression in liver. Low expression in colon, kidney, testis and pancreas. No expression in brain | liver: 2359.8 | Cluster 83: Liver - Metabolism |
| 9 | P19013 | 0.00096 | Keratin, type II cytoskeletal 4 | Epithelial | esophagus: 19104.8 | Cluster 29: Squamous epithelium - Innate immune response |
| 10 | P07101 | 0.00087 | Tyrosine 3-monooxygenase | Brain and adrenal gland | adrenal gland: 190.6;brain: 130.3 | Cluster 49: Pituitary gland - Hormone signaling |
| 11 | P09228 | 0.00085 | Cystatin-SA | Parotid gland and saliva | salivary gland: 5804.2 | Cluster 38: Salivary gland - Salivary secretion |
| 12 | P07196 | 0.00068 | Neurofilament light polypeptide | nan | brain: 183.3;retina: 86.8 | Cluster 17: Neuronal - Synaptic function |
| 13 | O95340 | 0.00067 | Bifunctional 3'-phosphoadenosine 5'-phosphosulfate synthase 2 | Cartilage and adrenal gland | adrenal gland: 131.0 | Cluster 11: Liver - Plasma proteins |
| 14 | P35558 | 0.00059 | Phosphoenolpyruvate carboxykinase, cytosolic [GTP] | Liver, kidney and adipocytes | kidney: 416.9;liver: 664.7 | Cluster 1: Liver & Kidney - Metabolism |
| 15 | P02760 | 0.00058 | Protein AMBP | Expressed by the liver, secreted in plasma, urine and CSF | liver: 10607.0 | Cluster 31: Liver - Plasma proteins |
| 16 | P14770 | 0.00053 | Platelet glycoprotein IX | nan | bone marrow: 5.0;lung: 3.1;lymphoid tissue: 8.7 | Cluster 81: Lymphoid tissue & Bone marrow - Innate immune response |
| 17 | Q8TDL5 | 0.00053 | BPI fold-containing family B member 1 | Epithelium of duodenum, trachea and lung | cervix: 546.4;salivary gland: 264.3;stomach 1: 499.1 | Cluster 75: Epithelium - Extracellular exosomes |
| 18 | P12036 | 0.00052 | Neurofilament heavy polypeptide | nan | brain: 95.6;prostate: 237.7 | Cluster 64: Neuronal - Transcription |
| 19 | O15230 | 0.00051 | Laminin subunit alpha-5 | Heart, lung, kidney, skeletal muscle, pancreas, retina and placenta |  | Cluster 66: Non-specific - Transcription |
| 20 | Q06520 | 0.00051 | Sulfotransferase 2A1 | Liver and adrenal gland | liver: 1043.0 | Cluster 83: Liver - Metabolism |

### Reuse of pan-cancer dataset (MSV000095036)

As MLMarker was trained exclusively on healthy, solid, tissue proteomes, this makes it particularly sensitive to proteomic changes introduced by disease states such as cancer. To assess the extent of this effect, we applied MLMarker to proteomic data from the Tüshaus *et al*. pan-cancer study^32^, which includes 1,220 FFPE tissue samples spanning multiple cancer types and several healthy control samples. We used the pre-processed MassIVE output files (MSV000095036) without re-analysing the raw data. The dataset covered a diverse range of cancers: colorectal cancer (CRC), diffuse large B-cell lymphoma (DLBCL), glioblastoma, HeLa cervical cancer, melanoma, pancreatic ductal adenocarcinoma (PDAC), and oral squamous cell carcinoma (OSCC).

The absence of cancer-specific proteomes, cell lines and certain tissues (e.g. skin, pancreas) within MLMarker training data, meant we had to map each cancer type to its closest corresponding healthy tissue and applied a penalty factor of 1. The HeLa cervical cancer cell line was excluded from absolute tissue mapping, as it no longer reflects native tissue but was considered later in comparative analyses.

To evaluate the advantage of the machine learning component of MLMarker to baseline atlas comparisons, we compared it to three reference atlases: (i) the MLMarker Training Atlas (the median tissue expression profiles used during model training), (ii) the Human Protein Atlas (HPA) antibody staining atlas, and (iii) ProteomicsDB (PDB) mass spectrometric atlas. For each atlas, we implemented two classification strategies: Spearman correlation and k-nearest neighbors (KNN) classification. We evaluated classification under two mapping schemes: (i) strict mapping, requiring exact one-to-one correspondence between cancer tissue of origin and predicted tissue; and (ii) extended mapping, allowing biologically related tissues to be considered correct predictions. Extended mappings were defined based on anatomical and functional relationships (e.g., colorectal cancer mapped to colon, small intestine, rectum, duodenum, or appendix; glioblastoma mapped to brain and pituitary gland; OSCC mapped to parotid gland and tonsil).

The classification accuracy of the highest predicted tissue (top-1 accuracy) (Fig 11) showed the optimal performance of MLMarker with 72.9% for extended mapping and 50.7% for strict mapping. The next-best performance, 42.9%, was observed for HPA using spearman correlation and extended mapping. The comparatively low accuracy of the MLMarker training further reinforces that the RF algorithm, rather than the underlying training data alone, drive the improved classification.

**Fig 11.**
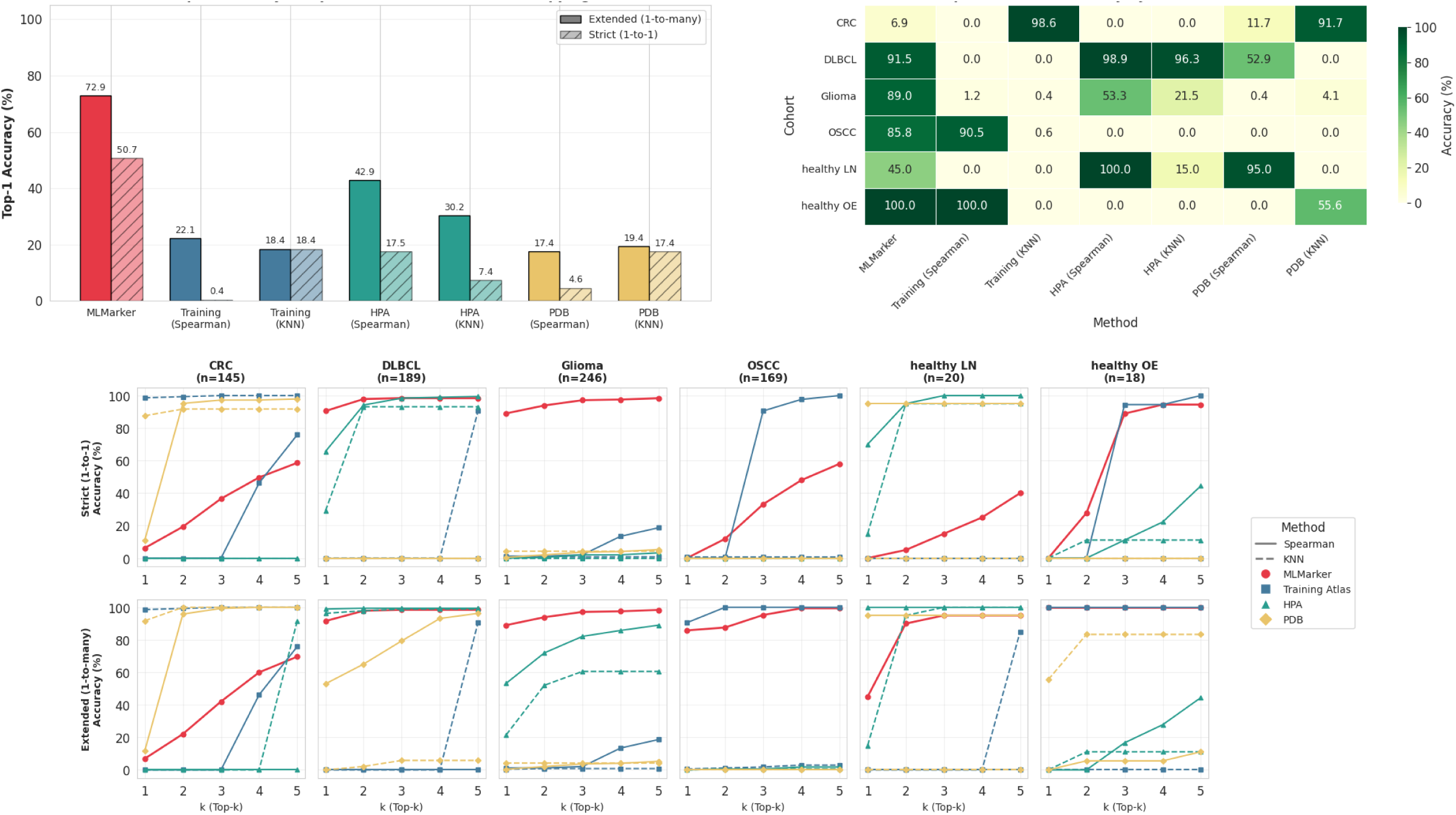
A) Top-1 accuracy (%) for strict (1-to-1) and extended (1-to-many) tissue mappings (y-axis) across classification methods (x-axis). MLMarker achieves the highest accuracy under both schemes, outperforming atlas-based Spearman and KNN approaches. B) Cohort-level accuracy heatmap showing classification performance (color scale, % accuracy) across cancer and healthy tissue cohorts (y-axis) for each method (x-axis). MLMarker maintains strong performance across most cohorts, with CRC emerging as the most challenging. C) Top-k accuracy (%) (y-axis) across k = 1–5 (x-axis) for each cohort, comparing MLMarker to atlas-based methods. MLMarker shows modest gains with increasing k, whereas atlas-based methods benefit substantially more, indicating that MLMarker’s top prediction is already highly informative.

Cohort level analyses show a strong deviation in classification performance. MLMarker maintained consistently high performance across most cancer types including DLBCL, glioma and OSCC but CRC emerged as a remarkably challenging cohort for most methods. Healthy cohorts (LN and OE) displayed a different pattern. Both showed strong MLMarker performance under extended mapping but substantially lower accuracy under strict mapping. This large gain indicates that these tissues share proteomic features with related anatomical sites, making extended mapping biologically appropriate. The same trend can be observed within the MLMarker Training Atlas pointing towards this being a training data limitation with limited tissue representation. These findings highlight the importance of expanding the training atlas to include underrepresented healthy tissues to further improve model generalizability.

The effect of varying the number of top predictions (k) also differed markedly between methods. Increasing k from 1 to 3 or 5 improved accuracy for all approaches, but the magnitude of improvement was substantially larger for atlas-based methods than for MLMarker (Fig 11). This pattern suggests that MLMarker’s top-ranked prediction is already highly informative and that the model captures finer-grained tissue-specific structure in the proteomic data. In contrast, atlas-based methods appear to rely more heavily on broader similarity patterns, benefiting disproportionately from the inclusion of additional candidate tissues. The relatively modest gains for MLMarker when expanding to top-5 predictions therefore reflect a higher degree of granularity and confidence in its primary classification.

To further validate these findings, we compared MLMarker’s predictions against two synthetic control samples: (i) a random sample, generated to mimic the original data distribution; and (ii) a zero-value sample to test the penalty factor performance. Both samples clustered distinctly from biological samples, reinforcing the biological relevance of MLMarker’s predictions and the efficiency of the penalty factor (Supplementary figure 7).

UMAP clustering of the original protein-level expression data revealed clear cancer-type-specific clusters, with some notable exceptions (Fig 12). Indeed, 30 melanoma samples are metastases from neuronal origin and clustered within the glioblastoma group, reflecting shared protein expression patterns. When restricting the UMAP to only use the MLMarker features, CRC and PDAC became more distinct, but melanoma samples remained interspersed within the glioblastoma cluster. At the prediction level, clustering became less clearly defined, with increased overlap between cancer types, mirroring the original study’s top 10 protein UMAP clustering. A striking exception was the HeLa cells dataset, which formed a dense, isolated cluster, indicating a highly distinct proteomic profile, overwhelmingly predicted as B-cells. HeLa cells, originating from cervical cancer tissue, are expected to be highly similar to other tissues in the MLMarker training data, such as endometrium, ovary, oviduct, and placenta, more than B-cells.

**Fig 12.**
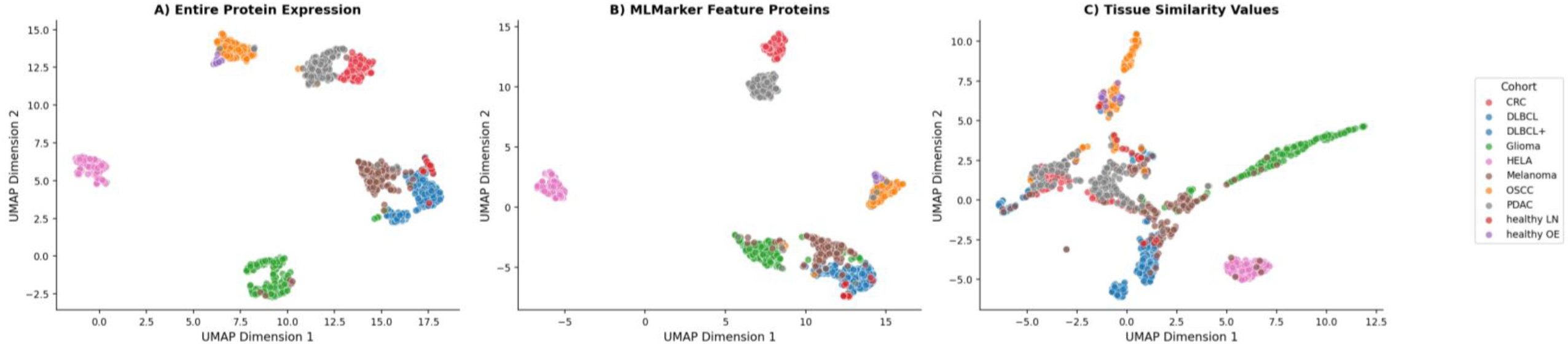
A). Dendrogram showing tissue similarity per cancer type, including random and fully zeroed samples (Random_Expression_1 and Zero_column respectively). B) UMAP protein expression of the entire dataset (top panel), protein expression of MLMarker features (middle panel) and tissue similarity values (bottom panel).

To explore this B-cell prediction, we first investigated the impact of missingness on the B-cell prediction by comparing the distribution of added protein features in HeLa samples with that of the entire dataset (Fig. 13a). This comparison revealed a skew towards higher missingness, though HeLa samples were not an outlier. Notably, B-cell similarity did not correlate directly with the number of added features (Fig 13b). We then examined whether the proteins contributing most to the B-cell similarity score were present or absent, given that missing features can also carry information. By separating proteins into present and absent groups and evaluating the direction of their SHAP values, we observed a similar distribution between both sets (Fig 13c). This indicates that the B-cell prediction is not solely driven by absent features. Instead, it suggests that the studied HeLa cell line possessed a proteomic profile more similar to B-cells than to other solid tissues in the reference set, possibly a result of immortalisation and extensive culturing.

**Fig 13.**
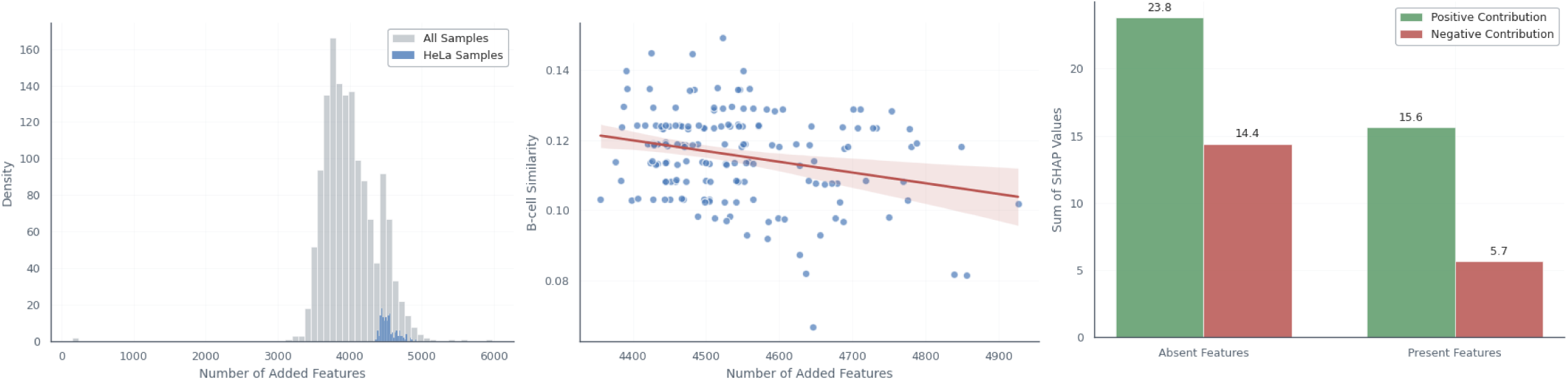
Analysis of B-cell similarity predictions in HeLa samples. A) Histogram showing the number of protein features added to complete the MLMarker feature space in HeLa samples compared to the full dataset. HeLa samples show a skew toward higher missingness but are not outliers. B) Scatterplot of B-cell similarity scores versus the number of added protein features in HeLa samples, showing no direct correlation. C) Distribution of SHAP value directions (positive or negative) for the top B-cell-contributing proteins, separated by presence or absence in the HeLa samples, indicating both present and absent features contribute similarly.

Glioblastoma and melanoma showed no distinct separate clustering. This is notable as MLMarker was never trained on skin, hence melanoma samples cannot be predicted correctly. However, MLMarker did have plentiful training data for brain tissue with some melanoma samples achieving very high brain similarity.

## Discussion

We here introduced MLMarker, a data-driven tissue prediction tool that leverages machine learning techniques to identify tissue-specific proteomic signatures, providing a new dimension to the analysis of complex biological data. In essence, MLMarker compares the presented proteome profile to a broad array of publicly available healthy tissue proteomes it was trained upon, providing a boosted DEA. In-depth analysis of the contributing proteins through SHAP analysis clarifies the model’s prediction process through Explainable AI and presents a tool for hypothesis generation. While originally designed to predict the tissue of samples, MLMarker’s ability to uncover underlying proteomic patterns offers a unique approach to generate hypotheses, making it a valuable tool not just for classification, but also for furthering biological understanding in various research contexts.

A prime example of MLMarker’s capability is illustrated in the study of cerebral melanoma metastases. The original study stratified tumours based on their response to treatment, but MLMarker presents a proteomic-based classification system, grouping samples based on their similarity to brain tissue. This shift in perspective leads to the hypothesis that certain melanoma tumours, especially those with poor treatment responses, might acquire proteomic characteristics that mimic brain tissue, facilitating their survival and adaptation in the brain’s microenvironment. This novel hypothesis was not captured with the original analyses and underscores how MLMarker can reveal new proteomic phenotypes. Following up this observation with differential expression analysis between the clear brain predicted and non-brain predicted sample led to a ten-fold increase in the number of differentially expressed proteins and therefore possibly biologically interesting targets.

By identifying brain-predicted samples, MLMarker revealed that these samples, regardless of their treatment response, retained a distinct set of brain-associated proteins. This opens up exploratory avenues for the study of these tumours adopting brain-like features through proteomic mimicry, enhancing their capacity to adapt to and survive within the brain environment. In contrast, non-brain-predicted tumours, particularly those with good responses to treatment, were enriched for immune-related pathways, reinforcing the idea that immune activation rather than a specific proteotype may be a key factor in successful treatment responses. These findings provide compelling evidence that tumours exhibiting proteomic similarities to brain tissue might possess an invasive phenotype, enabling them to infiltrate the brain and evade immune surveillance. This hypothesis is further supported by the detection of ten known invasion markers found exclusively in brain-predicted samples and absent in non-brain-predicted tumours. This distinction emphasizes the connection between proteomic profiles and the invasive potential of tumours.

In addition to its role in identifying proteomic features associated with tumour invasiveness, MLMarker reinforces the concept that cancer proteomes strongly diverge from their healthy counterpart^27,29^ and some deviate more than others. An unexpected finding in the pan-cancer study was the clustering of melanoma samples with glioblastoma, indicating that certain melanoma cases displayed proteomic patterns similar to those of glioblastoma. Although some of these were known neuronal metastases, this was not the case for each melanoma and requires further investigation into the specific proteins responsible for this observation. The pan-cancer analysis also highlighted that healthy lymph node (LN) and olfactory epithelium (OE) samples benefited strongly from extended tissue mappings. These tissues are underrepresented in the MLMarker training atlas, and their improved performance under extended mapping suggests that expanding the training dataset with additional healthy LN and OE proteomes would further enhance model accuracy.

One of the most valuable features of MLMarker is its ability to handle sparse or noisy data, an issue that often arises in proteomics studies. By incorporating a penalty factor and adjusting tissue mappings, MLMarker was able to improve classification accuracy and sensitivity, even when faced with missing or incomplete protein data. This capability is particularly relevant for clinical applications, where liquid biopsies and other biofluid samples are increasingly being used for cancer detection and monitoring. These samples, by nature, contain a higher degree of missing data compared to traditional tissue biopsies, making MLMarker’s robustness in handling incomplete data a significant advantage. In fact, MLMarker was able to identify the tissue of origin for tissue-leakage proteins (TLPs) in biofluids, offering a new approach for detecting and monitoring cancer through biofluid analysis. Interestingly, MLMarker also detects a notable presence of liver-associated proteins, a phenomenon previously observed in both brain tissue and CSF^41^. However, rather than misclassifying these samples, MLMarker correctly recognizes that the co-occurrence of brain proteins with these liver proteins still indicates a neuronal origin rather than hepatic contamination. This further underscores MLMarker’s ability to handle complex tissue compositions and its adaptability to sample types beyond those in its training data, provided that appropriate adjustments, such as penalty factors, are applied.

The results also clearly demonstrate when the penalty factor should be enabled. Sparse samples (biofluids, organoids, low-input samples such as single cells) and samples deviating from the training data (cancers) consistently benefit from the usage of the penalty factor providing stable rankings and preventing spurious tissue assignments. In contrast, dense tissues with high proteome coverage show only marginal gains and may experience mild overcorrection at low dropout levels. Therefore, we recommend enabling the penalty factor by default for low-coverage or heterogeneous samples, while applying it more cautiously for dense, high-quality tissue proteomes.

Furthermore, we do not recommend using imputation for MLMarker input datasets. Imputation methods attempt to infer missing protein abundances from observed patterns, yet MLMarker relies on the observed presence or absence of proteins to compare to its healthy reference atlas. Consequently, imputation risks fabricating biologically unsupported evidence. The penalty factor provides a principled alternative because it does not invent abundance values but instead moderates how much missing features can influence the tissue ranking and its SHAP-based explanation, thereby preserving interpretability while acknowledging uncertainty.

A remarkable observation during the analyses was the separate clustering of HeLa cells from both healthy and cancerous tissues, reflecting a distinct proteomic signature. Despite their cervical origin, these samples were not predicted as any possibly related tissues. Instead, they were consistently predicted as B-cells, raising the question of whether the HeLa proteome has diverged too far from its original tissue context, resulting in assignment to the nearest isolated cell class. This opens future opportunities to study cell line proteome profiles from a tissue point of view.

By leveraging a fine-grained Random Forest model, MLMarker does not force overly confident classifications but instead allows for some degree of ambiguity in its predictions. More broadly, MLMarker’s performance relative to atlas-based Spearman and kNN baselines highlights what is gained by using a machine-learning model. The Random Forest captures non-linear interactions between proteins that are invisible to correlation-based or centroid-based methods. As a result, the expected tissue may not always rank as the top prediction, reflecting the inherent complexity and overlap in proteomic profiles across tissues. Rather than being a limitation, this characteristic provides a biologically meaningful perspective, where examining the top n predicted tissues can reveal underlying relationships between sample proteomes and multiple tissue types, offering deeper insights into tissue similarity and potential cross-tissue influences.

Looking to the future, through expanding the training data with more healthy tissues as well as biofluids, we aim to increase the models’ applicability. Currently available as a Python package and a Streamlit app, MLMarker is designed to bridge the gap between wet-lab and dry-lab scientists. This makes it a versatile tool that can be used by a wide range of researchers, from experimental biologists to computational scientists, fostering collaboration across disciplines. However, as with any tool, users must be aware of its limitations. To aid with this, comprehensive documentation has been provided, which offers further context and guidance on using the tool effectively.

In the realm of clinical research, MLMarker holds significant promise for identifying tissue origins of metastases. The tool’s capacity to classify tissues based on their proteomic profiles rather than conventional treatment-response criteria opens up new avenues for understanding metastatic behaviour. Additionally, MLMarker could play a pivotal role in overcoming challenges in single-cell proteomics, where hypothesis generation is often limited. By replacing traditional clustering-based differential expression analysis (DEA) with MLMarker’s orthogonal classification method, the tool could avoid the circularity issue known as double dipping, in which the same clusters are used for both generating hypotheses and performing subsequent analyses. This could streamline the workflow and lead to more accurate, unbiased results in proteomics studies.

Ongoing improvements to MLMarker aim to address challenges related to missing values and further expand the dataset used in its training. However, integrating more public data remains challenging due to the continuing lack of comprehensive metadata in publicly available proteomics datasets. The importance of metadata cannot be overstated, as it is essential for ensuring reproducibility and enabling the reuse of datasets across research projects. The development of metadata standards, such as the SDRF-Proteomics metadata standard and tools like lesSDRF^42^ are critical steps forward in addressing this issue. Widespread adoption of these standards will not only improve data quality and accessibility but also accelerate the development of data-driven methods like MLMarker.

In conclusion, MLMarker represents a significant innovation in the field of proteomics, providing a data-driven tool for hypothesis generation and improving the interpretation of complex proteomic data. Its abilities to handle missing data, to classify tissues based on proteomic profiles, and to generate new biological insights make it a versatile and indispensable tool in both basic research and clinical applications. As MLMarker continues to evolve, it will undoubtedly contribute to a deeper understanding of cancer biology and other complex diseases, paving the way for more precise and personalized therapeutic strategies. The continued development and adoption of standards for proteomics data will further enhance the tool’s impact, ensuring that it remains a key resource for researchers seeking to unlock the full potential of proteomics in biomedical science.

## Conclusion

MLMarker moves beyond conventional classification and differential analysis by introducing a continuous, probabilistic framework for assessing tissue similarity in proteomics data. Instead of relying on predefined groups, the tool compares each sample against a reference atlas of healthy tissue proteomes, producing interpretable similarity scores that reflect underlying biology. The model operates within a fixed protein feature space and was trained on normalized expression data from 34 healthy tissues. A penalty factor corrects for the impact of missing proteins, making MLMarker robust to sparsity and suitable for complex samples such as biofluids or tumour tissues with partial proteome coverage. SHAP values provide a transparent link between input features and predicted tissue similarities, enabling the identification of proteins that drive classification.

MLMarker was particularly effective in creating new biological hypotheses. In cerebral melanoma metastases, MLMarker identified a subset of tumours with high similarity to brain tissue, pointing to proteomic adaptation to the brain microenvironment, an insight not captured by conventional analysis. The application on biofluid and cancer data, very distinct from the original healthy tissue training data demonstrated the model’s robustness and innovation. Importantly, MLMarker outperformed atlas-based Spearman and kNN baselines, reflecting its ability to capture non-linear structure and subtle tissue relationships that median tissue profiles cannot represent.

As proteomics expands toward lower-input samples and broader clinical use, MLMarker provides an interpretable, data-resilient tool for tissue inference and hypothesis generation across diverse experimental contexts.

## Materials and methods

### Public datasets used for MLMarker application

#### Cerebral Melanoma Metastases Proteomics (PXD007592)

The dataset PXD007592 from Zila *et al*., initially generated to investigate proteomic differences between good and poor responders to MAP kinase inhibitors (MAPKi) in cerebral melanoma metastases, was used in this study. Raw mass spectrometry data files were downloaded from the PRIDE repository and reprocessed using an updated computational workflow (outlined below in the Reprocessing Pipeline section). Sample metadata, including the treatment response classification (good vs. poor responders), was inferred directly from the file names provided in the original dataset.

#### Liver dataset (PXD009021)

The dataset PXD009021 from Davis et al. was a human liver tissue harvested from donors to establish an in-house used reference sample. Raw mass spectrometry data files were downloaded from the PRIDE repository and reprocessed using an updated computational workflow (outlined below in the Reprocessing Pipeline section)

#### Cerebrospinal Fluid (CSF) Proteome Dataset (PXD008029)

To assess the applicability of MLMarker to biofluid proteomics, cerebrospinal fluid (CSF) samples from the Yamana dataset PXD008029 were selected for analysis. Sample metadata was extracted from the file names provided in the original dataset.

#### Pan-Cancer Formalin-Fixed Paraffin-Embedded (FFPE) Proteomics Dataset (MSV000095036)

The dataset from Tüshaus *et al*. consists of proteomic profiles for 1,220 FFPE tumor samples from multiple cancer types. Unlike the other datasets, raw data were not reprocessed for this study. Instead, processed output files from the original study, which are publicly available on MassIVE, were used directly for MLMarker analysis. Sample metadata, including cancer type and tumor classification, was retrieved from the mapping file provided by the original study authors in the MassIVE files.

### Reprocessing pipeline

For datasets requiring reprocessing, specifically PXD007592 (melanoma) and PXD008029 (CSF), a standardized computational workflow was applied to ensure consistency across datasets.

#### Protein Identification

Ionbot^35^ (v0.11.0) was employed for open modification searches. The Uniprot human reference proteome (reviewed) was used for sequence matching. Carbamidomethylation of cysteine (C+57.021464 Da) was set as a fixed modification, while methionine oxidation (M+15.994915 Da) was set as a variable modification. The precursor mass tolerance was ±10 ppm, and the fragment mass tolerance was ±20 ppm. Peptide-spectrum match (PSM) false discovery rate (FDR) was controlled to be < 1%.

#### Protein Quantification

FlashLFQ ^36^(v1.0.3) was used for label-free quantification. The “match-between-runs” option was disabled to ensure independent quantification per sample.

#### Differential Expression Analysis

MSqRob2^37^(v1.4.0) was used for robust statistical analysis of differential expression. Peptide intensities were log2-transformed, median-centered for normalization, and aggregated to protein-level using robust summarization. Proteins detected in fewer than four samples were excluded. Statistical significance was determined using linear mixed models with Benjamini-Hochberg correction (α = 0.01). Pairwise contrasts were tested for: (i) good vs. poor responders, (ii) brain-predicted vs. non-brain-predicted samples, and (iii) hierarchical clustering-derived groups.

### The MLMarker package

MLMarker is implemented as a modular Python package that bundles a pretrained Random Forest classifier^26^ together with its expected feature list and a dedicated explainability component. An MLMarker class couples model loading, input validation, Normalizatoin, prediction, and post hoc interpretation.

#### Input format and feature validation

Samples are provided as a dataframe in which columns correspond to protein identifiers and rows to samples. Prior to inference, MLMarker applies deterministic feature validation to ensure compatibility with the fixed feature space.

This procedure retains only proteins present in the model specification, resolves duplicated columns, appends missing model features as zero abundance columns, enforces the exact model feature order, and returns a validated fixed length feature matrix.

#### Normalization

After validation, MLMarker applies per sample min max normalization. Each sample vector is rescaled using its within sample minimum and maximum, which reduces sensitivity to study specific abundance scales and emphasises relative abundance patterns within each sample. The resulting normalised matrix is used for all downstream prediction and interpretation.

#### Prediction and SHAP based score reconstruction

MLMarker produces a ranked list of tissues by applying the random forest model to the normalised feature matrix. For interpretability, SHAP values are computed using shap.TreeExplainer to obtain class specific feature attributions. For each predicted tissue class, MLMarker reconstructs the tissue score as the class specific expected value returned by the explainer plus the sum of per protein SHAP contributions, and tissues are ranked by these reconstructed scores. Predictions are returned as the top 𝑘tissues, enabling evaluation under both strict rank one criteria and more permissive rank based criteria when tissue mapping granularity differs across annotation schemes.

#### Penalty factor for missing feature effects

Because MLMarker operates in a fixed feature space, proteins not detected in a query sample are encoded as zero abundance. To limit the impact of ambiguous zeros on the SHAP decomposition and the resulting tissue ranking, MLMarker implements an optional penalty factor 𝜆that adjusts SHAP values for absent proteins. Absent proteins are defined as validated features with zero abundance in the query sample. During initialisation, MLMarker computes a zero input SHAP reference by evaluating the explainer on a feature vector in which all proteins are set to zero. During inference, if 𝜆is greater than zero, MLMarker identifies absent proteins that nevertheless contribute non zero SHAP mass across the selected tissues and subtracts 𝜆times their corresponding zero input SHAP reference contribution. SHAP values for present proteins and for absent proteins with zero SHAP contribution remain unchanged, and the adjusted SHAP table is returned in the original feature order.

#### Annotation utilities and visualisation

MLMarker provides optional utilities to map high contribution proteins to external knowledge sources and to summarise results visually. Protein annotations can be retrieved from UniProt, tissue expression summaries can be queried from the Human Protein Atlas, and functional enrichment can be performed with g:Profiler on protein sets derived from SHAP contributions. Visual outputs include tissue score summaries and per tissue contribution views. When spectral counts are used as input, MLMarker provides a Normalized Spectral Abundance Factor utility that normalises counts by protein length prior to prediction.

### Penalty factor evaluation by simulated protein missingness

Simulations were performed on two public datasets representing contrasting proteome depths namely PXD009021 (liver tissue) and PXD008029 (cerebrospinal fluid). For each dataset, the target tissue label used for evaluation was liver (PXD009021) and pituitary gland (PXD008029). Raw data were reprocessed using ionbot for peptide identification and FlashLFQ for protein quantification. Protein abundances were expressed as Normalized Spectral Abundance Factor (NSAF) values when required for abundance dependent missingness.

#### Missingness mechanisms

Protein dropout was simulated using three mechanisms that reflect common regimes in proteomics.

1. Missing completely at random MCAR Proteins were removed uniformly at random from the detected set, independent of abundance.
2. Missing not at random MNAR Proteins were removed with higher probability at lower abundance to emulate detection limit effects. Removal weights were derived from normalised NSAF values, using a severity parameter 𝜅 = 2, and proteins were sampled for removal in proportion to these weights.
3. Targeted missingness To emulate preferential loss of diagnostic proteins, a tissue informative marker set was defined as the top 50 proteins ranked by absolute SHAP importance for the target tissue. Marker proteins were assigned a fivefold higher removal weight than non markers, and proteins were sampled for removal in proportion to these weights.

#### Dropout grid and replication

Dropout was simulated at nine levels, 0 to 80 percent in 10 percent increments. For each level, the number of removed proteins was set by multiplying the dropout fraction by the number of proteins detected in the original sample and taking the integer floor. Each condition was repeated 10 times using distinct random seeds. Across datasets, missingness mechanisms, dropout levels, penalty strategies, and replicates, this yielded 3,240 simulation runs.

#### Penalty strategies

We evaluated fixed and adaptive penalty strategies.

1. Fixed penalties: A constant 𝜆 was applied to all samples, using 𝜆 = 0(no penalty), 𝜆 = 0.5(conservative), 𝜆 = 1(full), and 𝜆 = 2(aggressive).
2. Adaptive penalties: Adaptive strategies set 𝜆 as a function of sample feature coverage, defined as the fraction of model features that are non zero in the validated sample. Two adaptive functions were used, namely a piecewise linear interpolation and a logistic interpolation, both increasing 𝜆 as coverage decreased. Reference coverages were set to 0.42 and 0.08, reflecting high and low coverage regimes observed in the training data, and the maximum penalty was set to 𝜆_max_ = 2.

#### Evaluation metrics

For each simulated sample, MLMarker produced a ranked list of tissue scores. Performance was quantified using top 1 accuracy, defined as the fraction of runs in which the target tissue ranked first. Rank stability was assessed by binning target ranks into four categories, rank 1, rank 2, rank 3, and rank 4 or worse. Overcorrection was assessed by comparing accuracy under a penalty strategy to the no penalty baseline and defining an overcorrection event as an accuracy decrease greater than 0.01.

### Baseline atlas comparisons and alternative classifiers

#### Reference atlases

We compared MLMarker to correlation based and distance based classification using three reference atlases. The HPA data were downloaded from https://www.proteinatlas.org/about/download (number 27, version 21.0) and included the protein expression values (normal_tissue.tsv) which contains ordinal expression values based on immunohistochemistry observations of tissue microarrays with antibodies. ProteomicsDB data (version 4.0) contained the normalized MS1–iBAQ intensities. Human Protein Atlas expression summaries were aggregated into a protein by tissue matrix by UniProt accession and tissue. ProteomicsDB tissue profiles were extracted as normalised intensities after excluding cancer and biofluid entries. The MLMarker training atlas was summarised by computing mean protein profiles per tissue across the training samples.

#### Alternative classification methods

Spearman correlation classification assigned each sample to the tissue with the highest Spearman rank correlation between the sample profile and the reference tissue profile, using only proteins present in both and requiring at least 10 shared proteins. K nearest neighbours classification used correlation distance to identify the nearest training samples for the MLMarker training atlas, and treated tissue level profiles as pseudo samples for Human Protein Atlas and ProteomicsDB.

#### Accuracy Metrics

Classification accuracy was evaluated using Top-k accuracy, measuring whether the true tissue of origin appeared among the top k predictions (k = 1, 2, 3, 4, 5). Accuracy was reported for both strict and extended tissue mappings.

#### Interpretation analyses

Uniform Manifold Approximation and Projection (UMAP) was applied to visualize the data using the umap package v0.1.1.

SHAP (SHapley Additive exPlanations) values were calculated using the shap package (v0.46.0) to interpret the contributions of individual proteins to tissue classification predictions. Positive SHAP values were associated with proteins that increased the likelihood of a given tissue classification, while negative SHAP values indicated proteins that decreased the likelihood of the classification.

Over-representation analysis was performed using the GProfiler tool (v1.0.0) to identify enriched biological processes, molecular functions, and cellular components. KEGG and Reactome pathways, as well as tissue-specific expression data from the Human Protein Atlas, were also incorporated into the analysis. Proteins ranked by SHAP feature importance were used as input for this analysis. Benjamini-Hochberg correction was used to obtain the adjusted p-value.

Correlation analysis between adjusted SHAP values and adjusted p-values was performed using Pearson correlation coefficient.

## Data and code Availability

All datasets used in this study are publicly available via ProteomeXchange (PXD007592, PXD009021, PXD008029) and MassIVE (MSV000095036). MLMarker predictions, differential expression results, and SHAP-derived protein lists are available in the Supplementary Data & Github repository for the manuscript analyses, pip package and streamlit application respectively github.com/TineClaeys/MLMarker-manuscript; github.com/TineClaeys/MLMarker; github.com/TineClaeys/MLMarker-streamlit. The streamlit application is available through mlmarker.streamlit.app.

## Acknowledgements

T.C. and L.M. acknowledge funding from the Research Foundation Flanders (FWO) [ G010023N, G028821N, 12A8W25N]. L.M. acknowledges funding from the Horizon Europe Projects BAXERNA 2.0 [101080544] and COMBINE [101191739], from the Ghent University Concerted Research Action [BOF21/GOA/033] and the CHIST-ERA project ODEEP-EU [G0GDV23N].

## Supplementary materials

**Fig S1.**
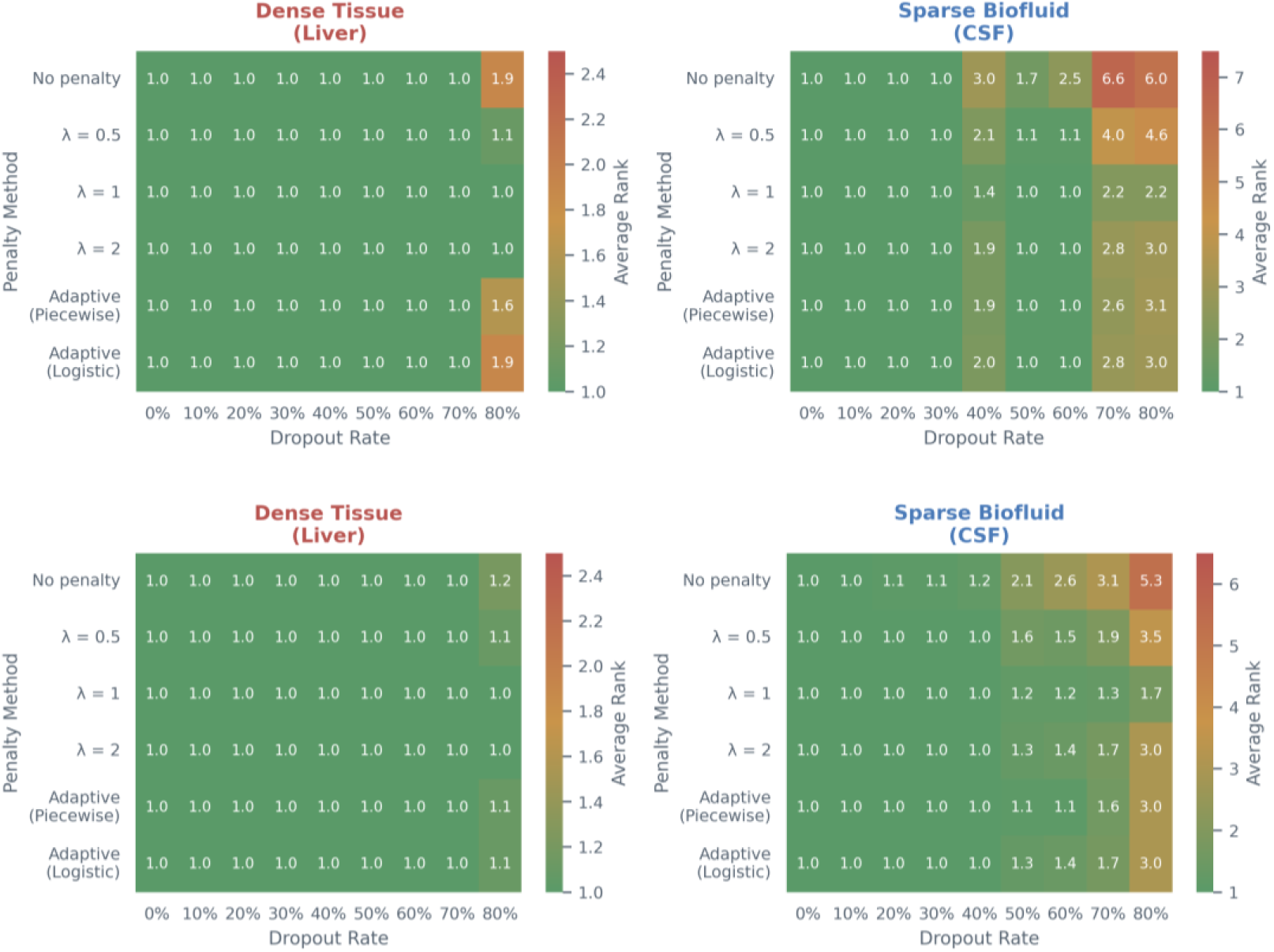
Average rank degradation of the correct tissue prediction under MCAR (top) and MNAR (bottom) missingness across penalty strategies for dense tissue (liver, left) and sparse biofluid (CSF, right). Optimal performance is indicated by a limited rank degradation. Each heatmap displays the average rank of the correct tissue prediction across increasing dropout rates (x-axis, 0–80%) for six penalty methods (y-axis): no penalty, fixed λ values (0.5, 1, 2), and adaptive strategies (piecewise, logistic). In both samples, λ = 1 consistently maintains low rank values across all dropout levels, outperforming other methods especially under high missingness

**Fig S2.**
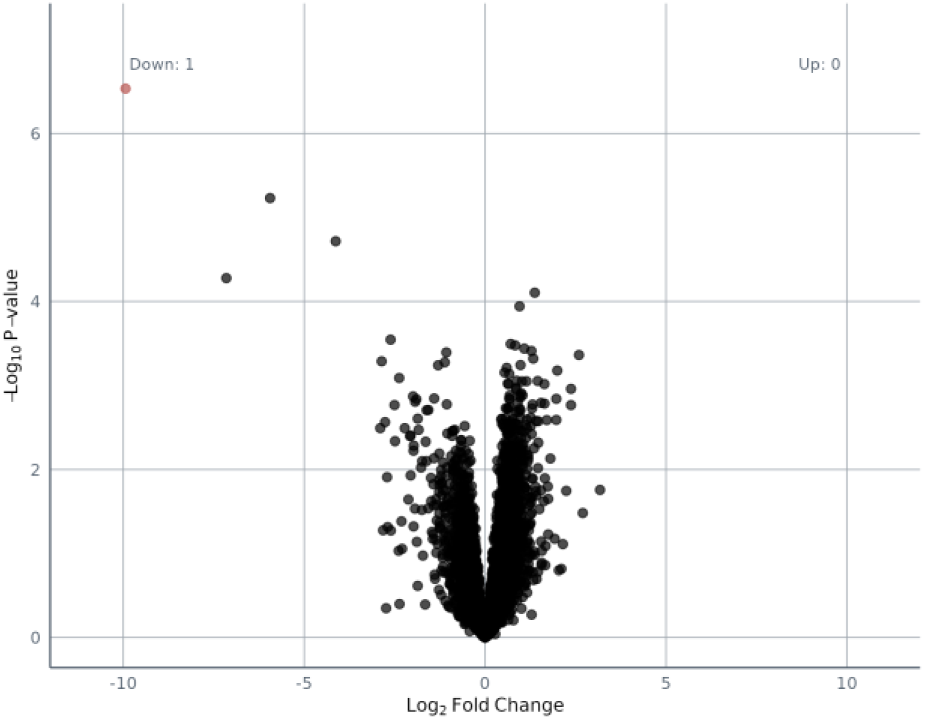
Differential expression analysis volcano plots between group A and group Bwith log fold change on X-axis, and -log_10_ adjusted p-value following Benjamini-Hochberg correction on Y-axis. Red dots indicate proteins with adjusted p-value < 0.01. Groups are previously determined through hierarchical clustering. Only 1 significantly downregulated protein in group B could be identified.

**Fig S3:**
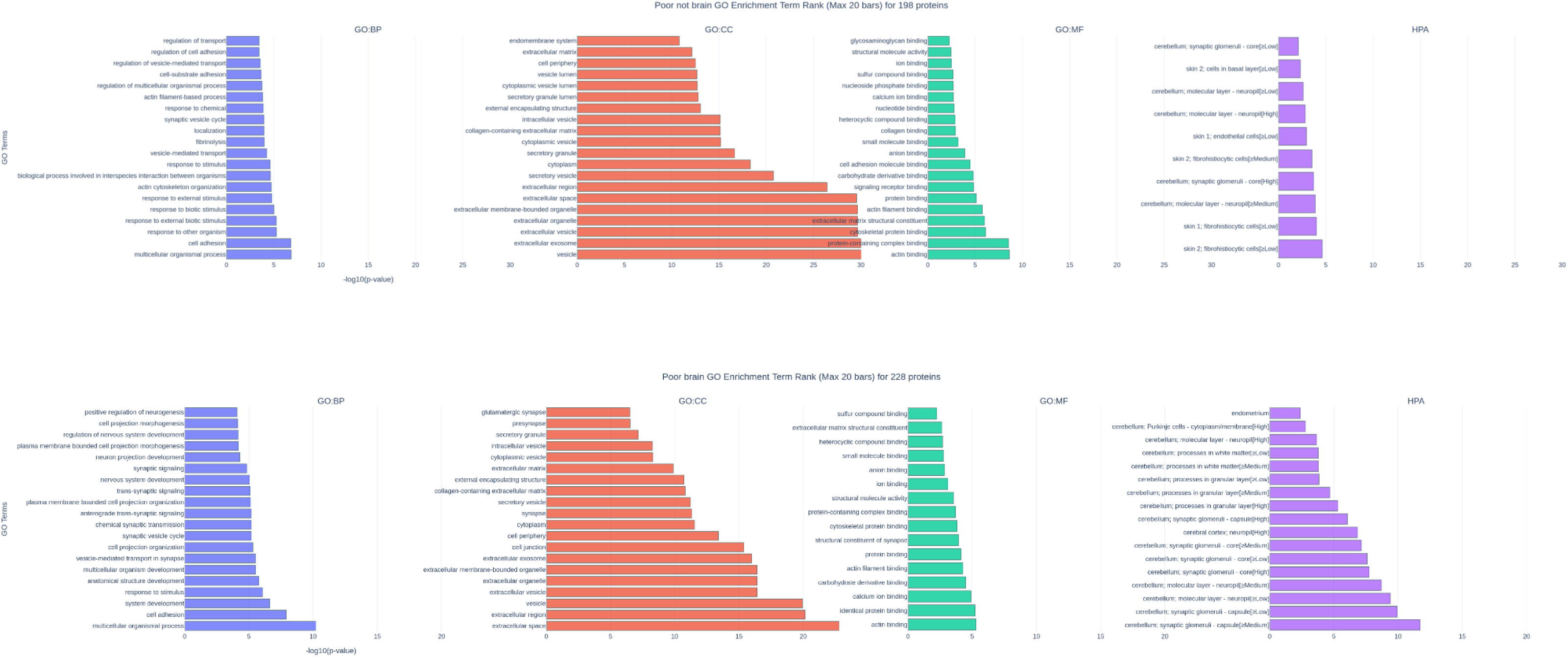
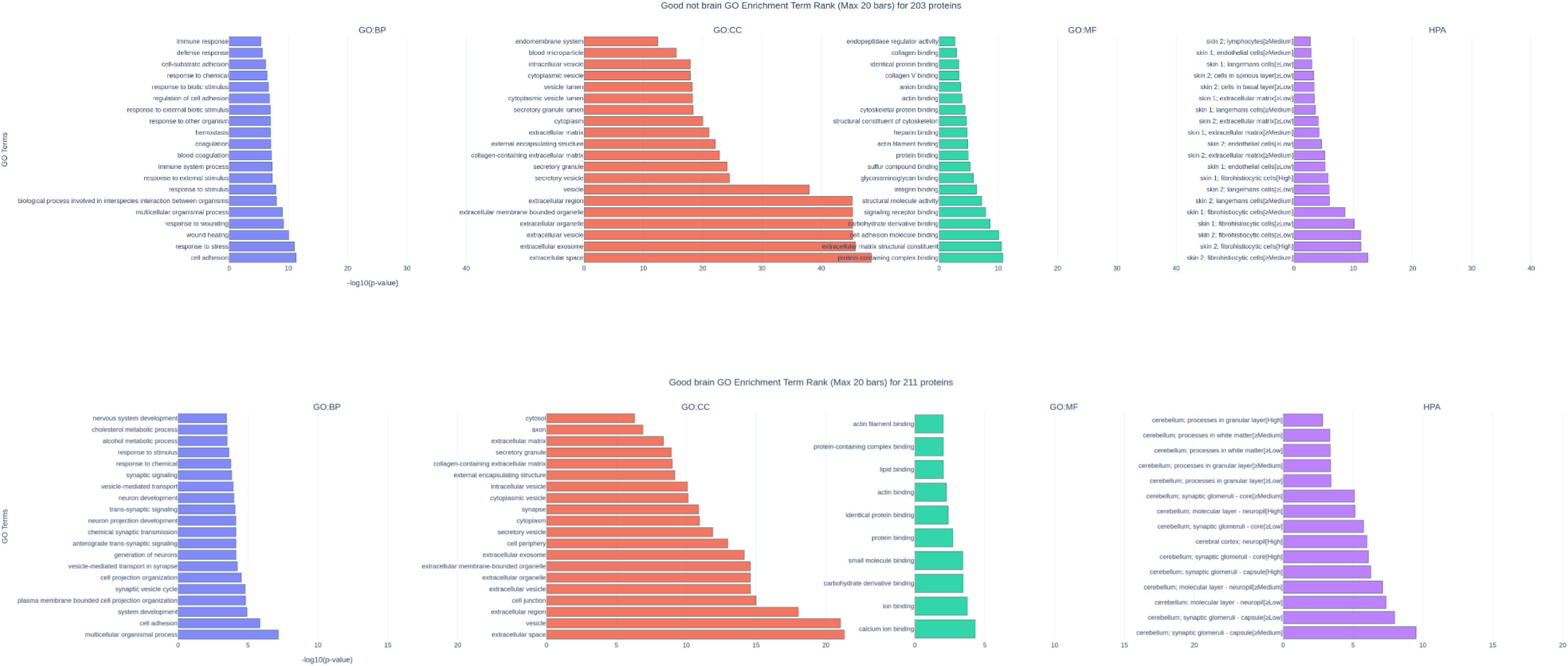
Over-representation analysis of the top 1% highest ranking proteins for the four subpopulations at p-value of 1%.

**Fig S4:**
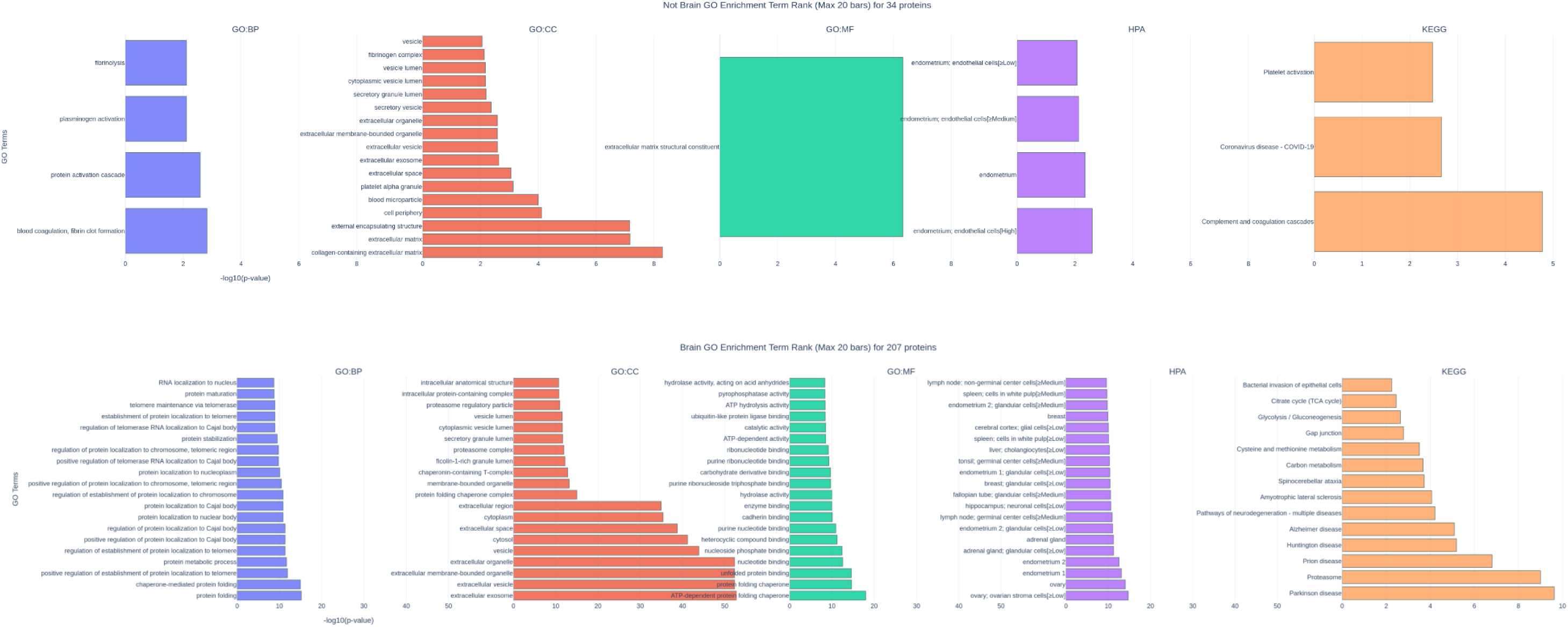
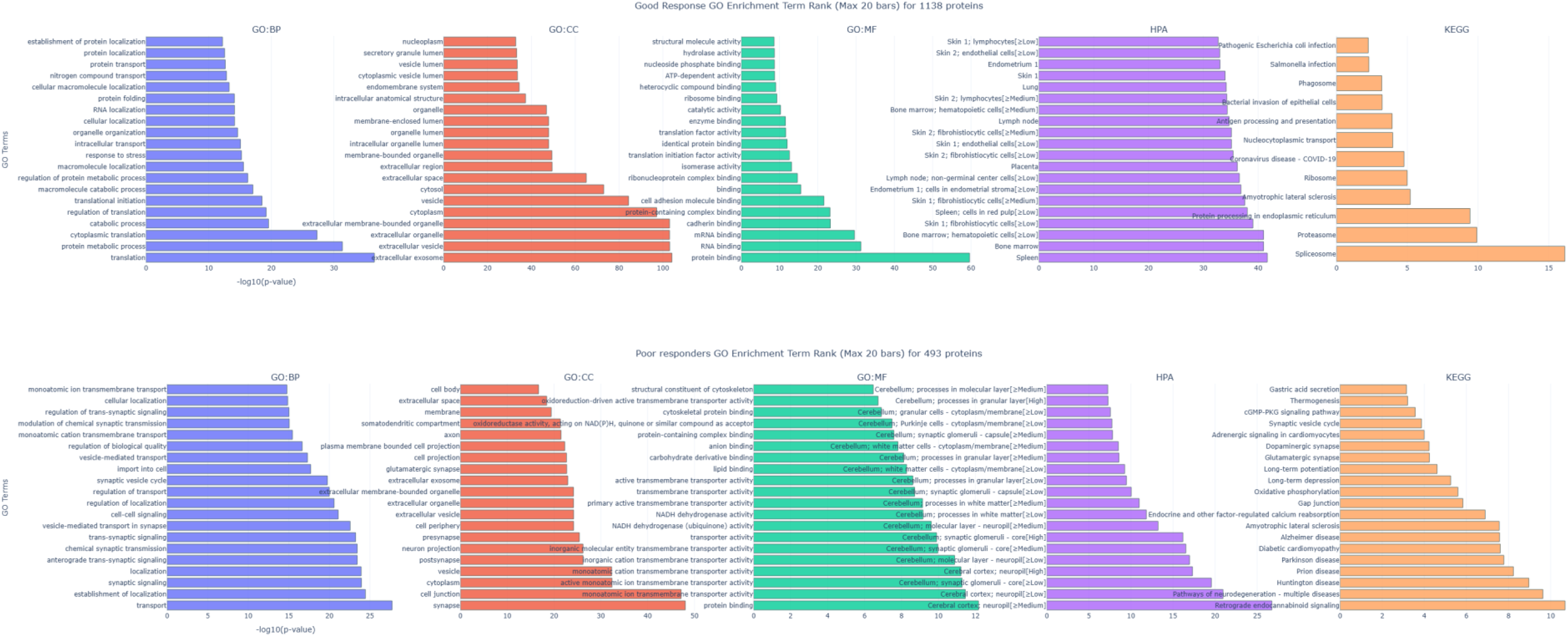
Over-representation analysis of the 255 differentially expressed proteins upregulated in the not-brain predicted samples (top) and brain predicted samples (bottom) at a p-value of 1 and of the differentially regulated proteins differentially regulated in good and poor responders.

**Fig S5:**
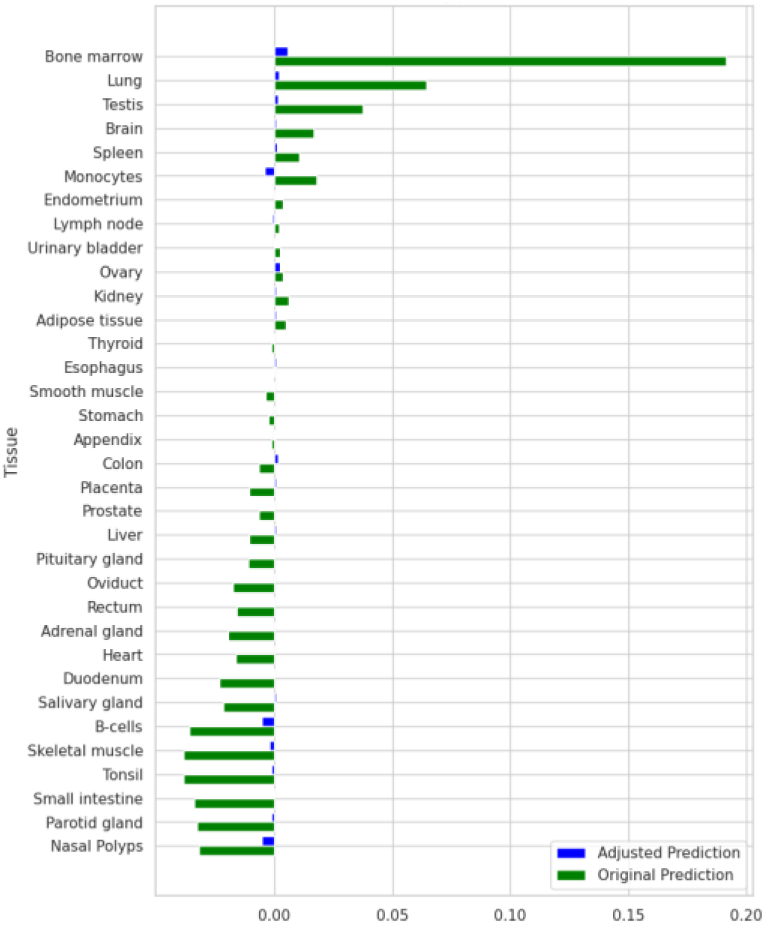
Bar plot showing tissue similarity scores for a fully zeroed sample. Original predictions are shown in green; adjusted predictions using the penalty factor are shown in blue. The application of the penalty factor results in a strong reduction in prediction confidence across all tissues.

**Fig S6:**
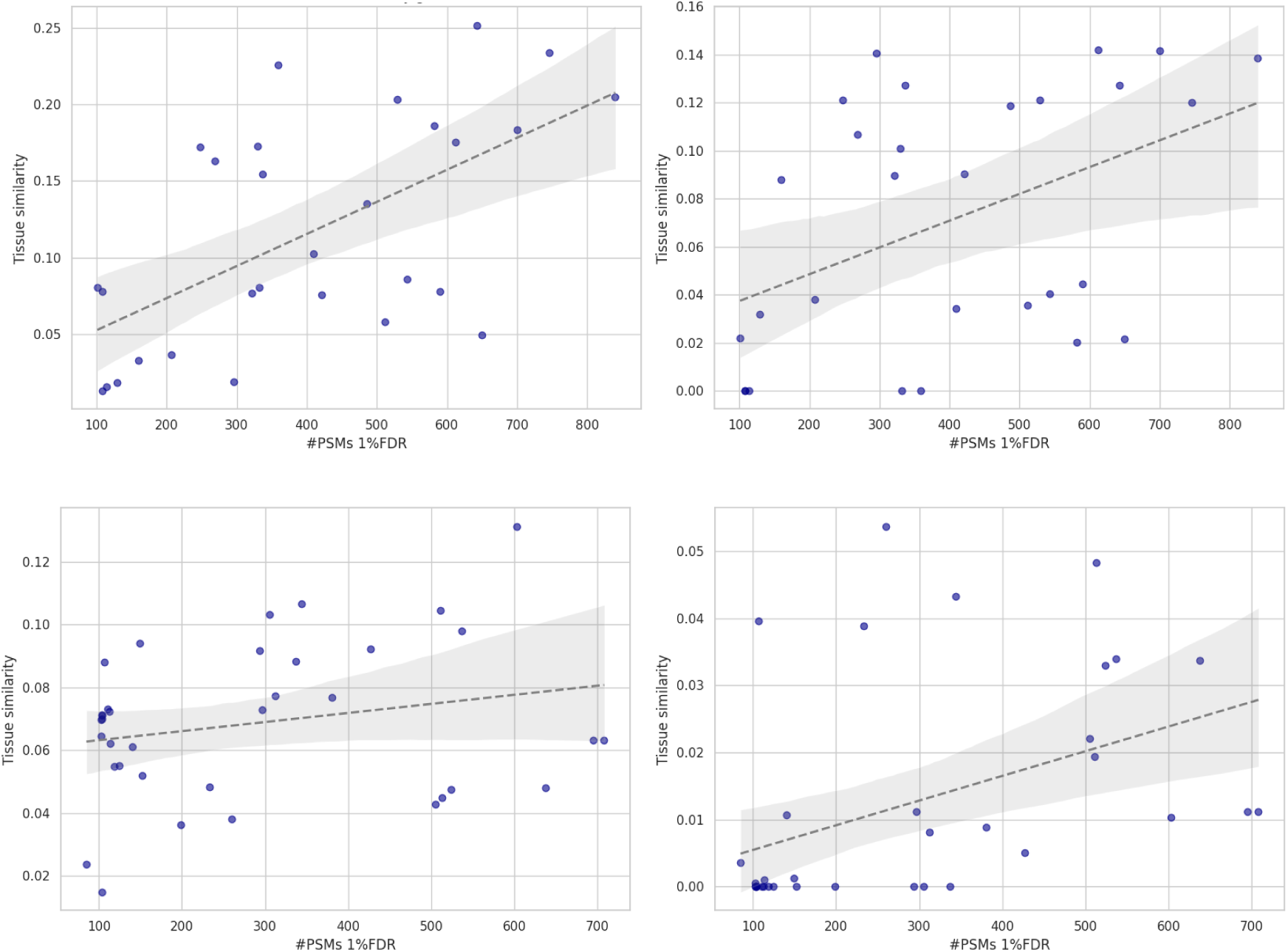
Scatterplot showing the relationship between tissue similarity and PSM counts for pituitary gland (top left), brain in the CSF samples (top right) and for adipose tissue (bottom left) and monocytes in plasma (bottom right). The x-axis represents the number of PSMs passing the 1% FDR threshold. A linear regression trendline indicates that an increase in PSMs often leads to greater certainty in tissue similarity predictions.

**Fig S7:**
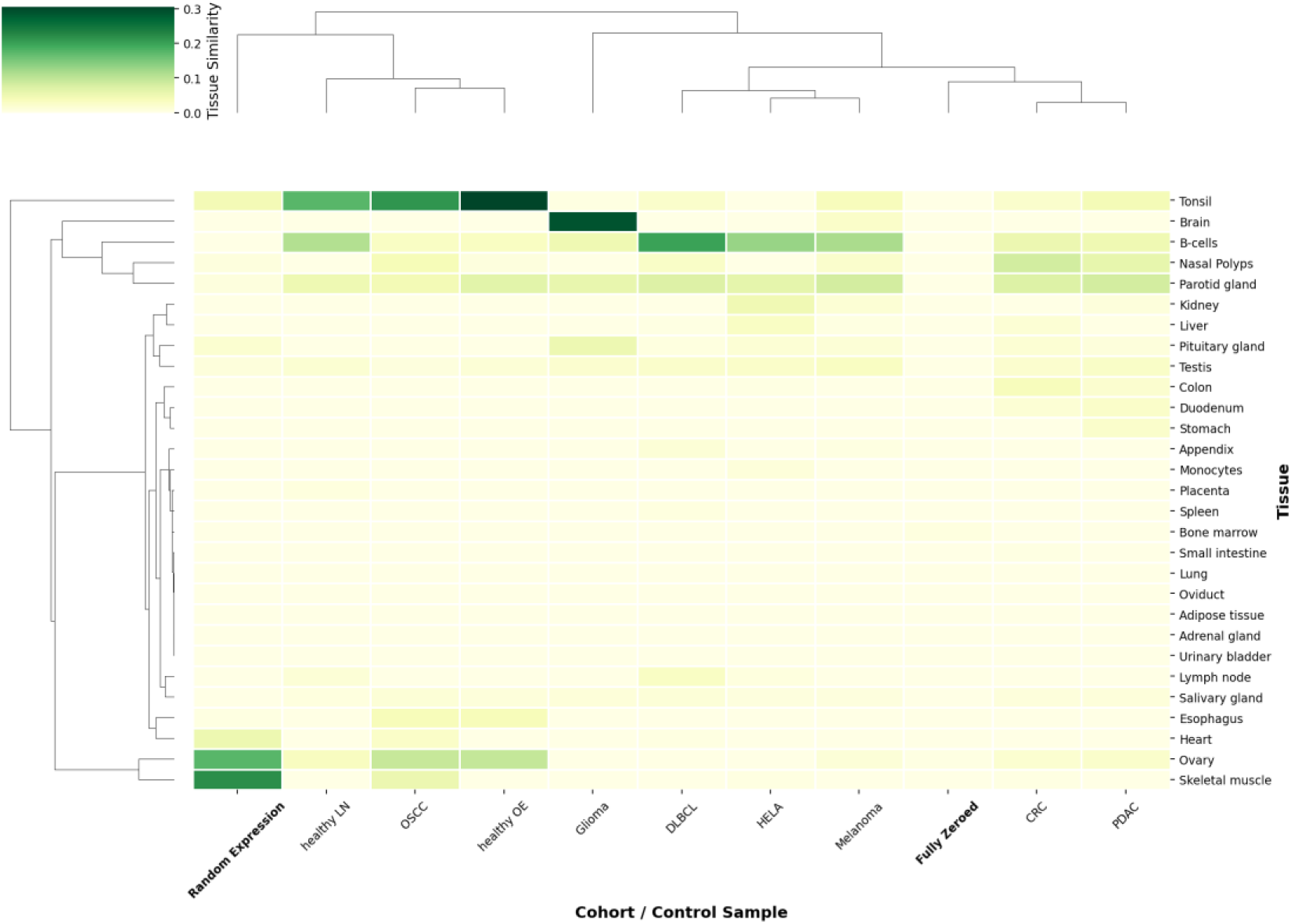
Dendrogram showing tissue similarity per cancer type, including random and fully zeroed samples (highlighted)

## Notes

### Competing Interest Statement

The authors have declared no competing interest.

### Summary of Updates

We added a new section titled Adaptability and explainability as core developments to make the methodological contributions explicit and to position MLMarker as a reusable framework rather than a proof of concept. Key technical updates include: (1) SHAP based reconstruction of tissue similarity scores, replacing direct random forest probabilities and enabling per protein interpretability; (2) a penalty factor framework to handle missing proteins, including a zero baseline SHAP reference and adaptive penalization strategies; (3) built in min max normalization to support heterogeneous quantification scales and acquisition modalities; (4) a fixed feature space alignment module to enable inference when protein overlap is incomplete, improving transferability; and (5) packaging MLMarker as a Python package with a Streamlit application to support user friendly deployment. We also added additional analyses examining intensity versus feature impact and benchmarking correlations with atlas based methods.

## References

(1) Krieger, J. R.; Wybenga-Groot, L. E.; Tong, J.; Bache, N.; Tsao, M. S.; Moran, M. F. Evosep One Enables Robust Deep Proteome Coverage Using Tandem Mass Tags While Significantly Reducing Instrument Time. J. Proteome Res. 2019, 18 (5), 2346–2353. 10.1021/acs.jproteome.9b00082.

(2) Ye, Z.; Sabatier, P.; van der Hoeven, L.; Lechner, M. Y.; Phlairaharn, T.; Guzman, U. H.; Liu, Z.; Huang, H.; Huang, M.; Li, X.; Hartlmayr, D.; Izaguirre, F.; Seth, A.; Joshi, H. J.; Rodin, S.; Grinnemo, K.-H.; Hørning, O. B.; Bekker-Jensen, D. B.; Bache, N.; Olsen, J. V. Enhanced Sensitivity and Scalability with a Chip-Tip Workflow Enables Deep Single-Cell Proteomics. Nat. Methods 2025, 1–11. 10.1038/s41592-024-02558-2.

(3) Wang, F.; Veth, T.; Kuipers, M.; Altelaar, M.; Stecker, K. E. Optimized Suspension Trapping Method for Phosphoproteomics Sample Preparation. Anal. Chem. 2023, 95 (25), 9471–9479. 10.1021/acs.analchem.3c00324.

(4) Ctortecka, C.; Clark, N. M.; Boyle, B. W.; Seth, A.; Mani, D. R.; Udeshi, N. D.; Carr, S. A. Automated Single-Cell Proteomics Providing Sufficient Proteome Depth to Study Complex Biology beyond Cell Type Classifications. Nat. Commun. 2024, 15 (1), 5707. 10.1038/s41467-024-49651-w.

(5) Bubis, J. A.; Arrey, T. N.; Damoc, E.; Delanghe, B.; Slovakova, J.; Sommer, T. M.; Kagawa, H.; Pichler, P.; Rivron, N.; Mechtler, K.; Matzinger, M. Challenging the Astral Mass Analyzer to Quantify up to 5,300 Proteins per Single Cell at Unseen Accuracy to Uncover Cellular Heterogeneity. Nat. Methods 2025, 1–10. 10.1038/s41592-024-02559-1.

(6) Stewart, H. I.; Grinfeld, D.; Giannakopulos, A.; Petzoldt, J.; Shanley, T.; Garland, M.; Denisov, E.; Peterson, A. C.; Damoc, E.; Zeller, M.; Arrey, T. N.; Pashkova, A.; Renuse, S.; Hakimi, A.; Kühn, A.; Biel, M.; Kreutzmann, A.; Hagedorn, B.; Colonius, I.; Schütz, A.; Stefes, A.; Dwivedi, A.; Mourad, D.; Hoek, M.; Reitemeier, B.; Cochems, P.; Kholomeev, A.; Ostermann, R.; Quiring, G.; Ochmann, M.; Möhring, S.; Wagner, A.; Petker, A.; Kanngiesser, S.; Wiedemeyer, M.; Balschun, W.; Hermanson, D.; Zabrouskov, V.; Makarov, A. A.; Hock, C. Parallelized Acquisition of Orbitrap and Astral Analyzers Enables High-Throughput Quantitative Analysis. Anal. Chem. 2023, 95 (42), 15656–15664. 10.1021/acs.analchem.3c02856.

(7) Declercq, A.; Bouwmeester, R.; Hirschler, A.; Carapito, C.; Degroeve, S.; Martens, L.; Gabriels, R. MS2Rescore: Data-Driven Rescoring Dramatically Boosts Immunopeptide Identification Rates. Mol. Cell. Proteomics MCP 2022, 21 (8), 100266. 10.1016/j.mcpro.2022.100266.

(8) Demichev, V.; Szyrwiel, L.; Yu, F.; Teo, G. C.; Rosenberger, G.; Niewienda, A.; Ludwig, D.; Decker, J.; Kaspar-Schoenefeld, S.; Lilley, K. S.; Mülleder, M.; Nesvizhskii, A. I.; Ralser, M. Dia-PASEF Data Analysis Using FragPipe and DIA-NN for Deep Proteomics of Low Sample Amounts. Nat. Commun. 2022, 13 (1), 3944. 10.1038/s41467-022-31492-0.

(9) Lazear, M. R. Sage: An Open-Source Tool for Fast Proteomics Searching and Quantification at Scale. J. Proteome Res. 2023, 22 (11), 3652–3659. 10.1021/acs.jproteome.3c00486.

(10) Teo, G. C.; Polasky, D. A.; Yu, F.; Nesvizhskii, A. I. Fast Deisotoping Algorithm and Its Implementation in the MSFragger Search Engine. J. Proteome Res. 2021, 20 (1), 498–505. 10.1021/acs.jproteome.0c00544.

(11) Domon, B.; Aebersold, R. Challenges and Opportunities in Proteomics Data Analysis*. Mol. Cell. Proteomics 2006, 5 (10), 1921–1926. 10.1074/mcp.R600012-MCP200.

(12) Aebersold, R.; Agar, J. N.; Amster, I. J.; Baker, M. S.; Bertozzi, C. R.; Boja, E. S.; Costello, C. E.; Cravatt, B. F.; Fenselau, C.; Garcia, B. A.; Ge, Y.; Gunawardena, J.; Hendrickson, R. C.; Hergenrother, P. J.; Huber, C. G.; Ivanov, A. R.; Jensen, O. N.; Jewett, M. C.; Kelleher, N. L.; Kiessling, L. L.; Krogan, N. J.; Larsen, M. R.; Loo, J. A.; Ogorzalek Loo, R. R.; Lundberg, E.; MacCoss, M. J.; Mallick, P.; Mootha, V. K.; Mrksich, M.; Muir, T. W.; Patrie, S. M.; Pesavento, J. J.; Pitteri, S. J.; Rodriguez, H.; Saghatelian, A.; Sandoval, W.; Schlüter, H.; Sechi, S.; Slavoff, S. A.; Smith, L. M.; Snyder, M. P.; Thomas, P. M.; Uhlén, M.; Van Eyk, J. E.; Vidal, M.; Walt, D. R.; White, F. M.; Williams, E. R.; Wohlschlager, T.; Wysocki, V. H.; Yates, N. A.; Young, N. L.; Zhang, B. How Many Human Proteoforms Are There? Nat. Chem. Biol. 2018, 14 (3), 206–214. 10.1038/nchembio.2576.

(13) Peng, H.; Wang, H.; Kong, W.; Li, J.; Goh, W. W. B. Optimizing Differential Expression Analysis for Proteomics Data via High-Performing Rules and Ensemble Inference. Nat. Commun. 2024, 15 (1), 3922. 10.1038/s41467-024-47899-w.

(14) Langley, S. R.; Mayr, M. Comparative Analysis of Statistical Methods Used for Detecting Differential Expression in Label-Free Mass Spectrometry Proteomics. J. Proteomics 2015, 129, 83–92. 10.1016/j.jprot.2015.07.012.

(15) Vizcaíno, J. A.; Deutsch, E. W.; Wang, R.; Csordas, A.; Reisinger, F.; Ríos, D.; Dianes, J. A.; Sun, Z.; Farrah, T.; Bandeira, N. ProteomeXchange Provides Globally Coordinated Proteomics Data Submission and Dissemination. Nat. Biotechnol. 2014, 32 (3), 223–226.

(16) Perez-Riverol, Y.; Bandla, C.; Kundu, D. J.; Kamatchinathan, S.; Bai, J.; Hewapathirana, S.; John, N. S.; Prakash, A.; Walzer, M.; Wang, S.; Vizcaíno, J. A. The PRIDE Database at 20 Years: 2025 Update. Nucleic Acids Res. 2025, 53 (D1), D543–D553. 10.1093/nar/gkae1011.

(17) Desiere, F.; Deutsch, E. W.; King, N. L.; Nesvizhskii, A. I.; Mallick, P.; Eng, J.; Chen, S.; Eddes, J.; Loevenich, S. N.; Aebersold, R. The PeptideAtlas Project. Nucleic Acids Res. 2006, 34 (suppl_1), D655–D658. 10.1093/nar/gkj040.

(18) Schmidt, T.; Samaras, P.; Frejno, M.; Gessulat, S.; Barnert, M.; Kienegger, H.; Krcmar, H.; Schlegl, J.; Ehrlich, H.-C.; Aiche, S.; Kuster, B.; Wilhelm, M. ProteomicsDB. Nucleic Acids Res. 2018, 46 (D1), D1271–D1281. 10.1093/nar/gkx1029.

(19) Ramasamy, P.; Turan, D.; Tichshenko, N.; Hulstaert, N.; Vandermarliere, E.; Vranken, W.; Martens, L. Scop3P: A Comprehensive Resource of Human Phosphosites within Their Full Context. J. Proteome Res. 2020, 19 (8), 3478–3486. 10.1021/acs.jproteome.0c00306.

(20) Degroeve, S.; Martens, L. MS2PIP: A Tool for MS/MS Peak Intensity Prediction. Bioinforma. Oxf. Engl. 2013, 29 (24), 3199–3203. 10.1093/bioinformatics/btt544.

(21) Bouwmeester, R.; Gabriels, R.; Hulstaert, N.; Martens, L.; Degroeve, S. DeepLC Can Predict Retention Times for Peptides That Carry As-yet Unseen Modifications. Nat. Methods 2021, 18 (11), 1363–1369. 10.1038/s41592-021-01301-5.

(22) Declercq, A.; Devreese, R.; Scheid, J.; Jachmann, C.; Van Den Bossche, T.; Preikschat, A.; Gomez-Zepeda, D.; Rijal, J. B.; Hirschler, A.; Krieger, J. R.; Srikumar, T.; Rosenberger, G.; Martelli, C.; Trede, D.; Carapito, C.; Tenzer, S.; Walz, J. S.; Degroeve, S.; Bouwmeester, R.; Martens, L.; Gabriels, R. TIMS2Rescore: A Data Dependent Acquisition-Parallel Accumulation and Serial Fragmentation-Optimized Data-Driven Rescoring Pipeline Based on MS2Rescore. J. Proteome Res. 2025, 24 (3), 1067–1076. 10.1021/acs.jproteome.4c00609.

(23) Robles, J.; Prakash, A.; Vizcaíno, J. A.; Casal, J. I. Integrated Meta-Analysis of Colorectal Cancer Public Proteomic Datasets for Biomarker Discovery and Validation. PLOS Comput. Biol. 2024, 20 (1), e1011828-.

(24) Camacho, O. J. M.; Ramsbottom, K. A.; Prakash, A.; Sun, Z.; Riverol, Y. P.; Bowler-Barnett, E.; Martin, M.; Fan, J.; Deutsch, E. W.; Vizcaíno, J. A.; Jones, A. R. Phosphorylation in the *Plasmodium Falciparum* Proteome: A Meta-Analysis of Publicly Available Data Sets. bioRxiv 2023, 2023.11.20.567785. 10.1101/2023.11.20.567785.

(25) Ochoa, D.; Jarnuczak, A. F.; Viéitez, C.; Gehre, M.; Soucheray, M.; Mateus, A.; Kleefeldt, A. A.; Hill, A.; Garcia-Alonso, L.; Stein, F.; Krogan, N. J.; Savitski, M. M.; Swaney, D. L.; Vizcaíno, J. A.; Noh, K.-M.; Beltrao, P. The Functional Landscape of the Human Phosphoproteome. Nat. Biotechnol. 2020, 38 (3), 365–373. 10.1038/s41587-019-0344-3.

(26) Claeys, T.; Menu, M.; Bouwmeester, R.; Gevaert, K.; Martens, L. Machine Learning on Large-Scale Proteomics Data Identifies Tissue and Cell-Type Specific Proteins. J. Proteome Res. 2023. 10.1021/acs.jproteome.2c00644.

(27) Kosti, I.; Jain, N.; Aran, D.; Butte, A. J.; Sirota, M. Cross-Tissue Analysis of Gene and Protein Expression in Normal and Cancer Tissues. Sci. Rep. 2016, 6, 24799. 10.1038/srep24799.

(28) De Wilde, J.; Van Paemel, R.; De Koker, A.; Roelandt, S.; Van de Velde, S.; Callewaert, N.; Van Dorpe, J.; Creytens, D.; De Wilde, B.; De Preter, K. A Fast, Affordable, and Minimally Invasive Diagnostic Test for Cancer of Unknown Primary Using DNA Methylation Profiling. Lab. Invest. 2024, 104 (8), 102091. 10.1016/j.labinv.2024.102091.

(29) Jiang, L.; Wang, M.; Lin, S.; Jian, R.; Li, X.; Chan, J.; Dong, G.; Fang, H.; Robinson, A. E.; Aguet, F.; Anand, S.; Ardlie, K. G.; Gabriel, S.; Getz, G.; Graubert, A.; Hadley, K.; Handsaker, R. E.; Huang, K. H.; Kashin, S.; MacArthur, D. G.; Meier, S. R.; Nedzel, J. L.; Nguyen, D. Y.; Segrè, A. V.; Todres, E.; Balliu, B.; Barbeira, A. N.; Battle, A.; Bonazzola, R.; Brown, A.; Brown, C. D.; Castel, S. E.; Conrad, D.; Cotter, D. J.; Cox, N.; Das, S.; de Goede, O. M.; Dermitzakis, E. T.; Engelhardt, B. E.; Eskin, E.; Eulalio, T. Y.; Ferraro, N. M.; Flynn, E.; Fresard, L.; Gamazon, E. R.; Garrido-Martín, D.; Gay, N. R.; Guigó, R.; Hamel, A. R.; He, Y.; Hoffman, P. J.; Hormozdiari, F.; Hou, L.; Im, H. K.; Jo, B.; Kasela, S.; Kellis, M.; Kim-Hellmuth, S.; Kwong, A.; Lappalainen, T.; Li, X.; Liang, Y.; Mangul, S.; Mohammadi, P.; Montgomery, S. B.; Muñoz-Aguirre, M.; Nachun, D. C.; Nobel, A. B.; Oliva, M.; Park, Y. S.; Park, Y.; Parsana, P.; Reverter, F.; Rouhana, J. M.; Sabatti, C.; Saha, A.; Skol, A. D.; Stephens, M.; Stranger, B. E.; Strober, B. J.; Teran, N. A.; Viñuela, A.; Wang, G.; Wen, X.; Wright, F.; Wucher, V.; Zou, Y.; Ferreira, P. G.; Li, G.; Melé, M.; Yeger-Lotem, E.; Barcus, M. E.; Bradbury, D.; Krubit, T.; McLean, J. A.; Qi, L.; Robinson, K.; Roche, N. V.; Smith, A. M.; Sobin, L.; Tabor, D. E.; Undale, A.; Bridge, J.; Brigham, L. E.; Foster, B. A.; Gillard, B. M.; Hasz, R.; Hunter, M.; Johns, C.; Johnson, M.; Karasik, E.; Kopen, G.; Leinweber, W. F.; McDonald, A.; Moser, M. T.; Myer, K.; Ramsey, K. D.; Roe, B.; Shad, S.; Thomas, J. A.; Walters, G.; Washington, M.; Wheeler, J.; Jewell, S. D.; Rohrer, D. C.; Valley, D. R.; Davis, D. A.; Mash, D. C.; Branton, P. A.; Barker, L. K.; Gardiner, H. M.; Mosavel, M.; Siminoff, L. A.; Flicek, P.; Haeussler, M.; Juettemann, T.; Kent, W. J.; Lee, C. M.; Powell, C. C.; Rosenbloom, K. R.; Ruffier, M.; Sheppard, D.; Taylor, K.; Trevanion, S. J.; Zerbino, D. R.; Abell, N. S.; Akey, J.; Chen, L.; Demanelis, K.; Doherty, J. A.; Feinberg, A. P.; Hansen, K. D.; Hickey, P. F.; Jasmine, F.; Kaul, R.; Kibriya, M. G.; Li, J. B.; Li, Q.; Linder, S. E.; Pierce, B. L.; Rizzardi, L. F.; Smith, K. S.; Stamatoyannopoulos, J.; Tang, H.; Carithers, L. J.; Guan, P.; Koester, S. E.; Little, A. R.; Moore, H. M.; Nierras, C. R.; Rao, A. K.; Vaught, J. B.; Volpi, S.; Snyder, M. P. A Quantitative Proteome Map of the Human Body. Cell 2020, 183 (1), 269–283.e19. 10.1016/j.cell.2020.08.036.

(30) Zila, N.; Bileck, A.; Muqaku, B.; Janker, L.; Eichhoff, O. M.; Cheng, P. F.; Dummer, R.; Levesque, M. P.; Gerner, C.; Paulitschke, V. Proteomics-Based Insights into Mitogen-Activated Protein Kinase Inhibitor Resistance of Cerebral Melanoma Metastases. Clin. Proteomics 2018, 15 (1), 13. 10.1186/s12014-018-9189-x.

(31) Rapid and Deep Profiling of Human Induced Pluripotent Stem Cell Proteome by One-Shot NanoLC–MS/MS Analysis with Meter-Scale Monolithic Silica Columns | Journal of Proteome Research.

(32) Tüshaus, J.; Eckert, S.; Schliemann, M.; Zhou, Y.; Pfeiffer, P.; Halves, C.; Fusco, F.; Weigel, J.; Hönikl, L.; Butenschön, V.; Todorova, R.; Rauert-Wunderlich, H.; The, M.; Rosenwald, A.; Heinemann, V.; Holch, J.; Steiger, K.; Delbridge, C.; Meyer, B.; Weichert, W.; Mogler, C.; Kuhn, P.-H.; Kuster, B. Towards Routine Proteome Profiling of FFPE Tissue: Insights from a 1,220-Case Pan-Cancer Study. EMBO J. 2025, 44 (1), 304–329. 10.1038/s44318-024-00289-w.

(33) Davis, W. C.; Kilpatrick, L. E.; Ellisor, D. L.; Neely, B. A. Characterization of a Human Liver Reference Material Fit for Proteomics Applications. Sci. Data 2019, 6, 324. 10.1038/s41597-019-0336-7.

(34) Yamana, R.; Iwasaki, M.; Wakabayashi, M.; Nakagawa, M.; Yamanaka, S.; Ishihama, Y. Rapid and Deep Profiling of Human Induced Pluripotent Stem Cell Proteome by One-Shot NanoLC-MS/MS Analysis with Meter-Scale Monolithic Silica Columns. J. Proteome Res. 2013, 12 (1), 214–221. 10.1021/pr300837u.

(35) Degroeve, S.; Gabriels, R.; Velghe, K.; Bouwmeester, R.; Tichshenko, N.; Martens, L. Ionbot: A Novel, Innovative and Sensitive Machine Learning Approach to LC-MS/MS Peptide Identification. bioRxiv 2022, 2021.07.02.450686. 10.1101/2021.07.02.450686.

(36) Millikin, R. J.; Solntsev, S. K.; Shortreed, M. R.; Smith, L. M. Ultrafast Peptide Label-Free Quantification with FlashLFQ. J. Proteome Res. 2018, 17 (1), 386–391. 10.1021/acs.jproteome.7b00608.

(37) Goeminne, L. J. E.; Sticker, A.; Martens, L.; Gevaert, K.; Clement, L. MSqRob Takes the Missing Hurdle: Uniting Intensity- and Count-Based Proteomics. Anal. Chem. 2020, 92 (9), 6278–6287. 10.1021/acs.analchem.9b04375.

(38) Nitz, A. A.; Giraldez Chavez, J. H.; Eliason, Z. G.; Payne, S. H. Are We There Yet? Assessing the Readiness of Single-Cell Proteomics to Answer Biological Hypotheses. J. Proteome Res. 2024, 24 (4), 1482–1492. 10.1021/acs.jproteome.4c00091.

(39) Pouliquen, D. L.; Boissard, A.; Coqueret, O.; Guette, C. Biomarkers of Tumor Invasiveness in Proteomics (Review). Int. J. Oncol. 2020, 57 (2), 409–432. 10.3892/ijo.2020.5075.

(40) Macron, C.; Lane, L.; Núñez Galindo, A.; Dayon, L. Identification of Missing Proteins in Normal Human Cerebrospinal Fluid. J. Proteome Res. 2018, 17 (12), 4315–4319. 10.1021/acs.jproteome.8b00194.

(41) Fang, Q.; Strand, A.; Law, W.; Faca, V. M.; Fitzgibbon, M. P.; Hamel, N.; Houle, B.; Liu, X.; May, D. H.; Poschmann, G.; Roy, L.; Stühler, K.; Ying, W.; Zhang, J.; Zheng, Z.; Bergeron, J. J. M.; Hanash, S.; He, F.; Leavitt, B. R.; Meyer, H. E.; Qian, X.; McIntosh, M. W. Brain-Specific Proteins Decline in the Cerebrospinal Fluid of Humans with Huntington Disease. Mol. Cell. Proteomics MCP 2009, 8 (3), 451–466. 10.1074/mcp.M800231-MCP200.

(42) Claeys, T.; Van Den Bossche, T.; Perez-Riverol, Y.; Gevaert, K.; Vizcaíno, J. A.; Martens, L. lesSDRF Is More: Maximizing The Value Of Proteomics Data Through Streamlined Metadata Annotation. Nat. Commun. 2023, 14 (6473). 10.21203/rs.3.rs-2937726/v1.

